# Neural support for contributions of utility and narrative processing of evidence in juror decision making

**DOI:** 10.1101/2020.11.11.378935

**Authors:** Jaime J. Castrellon, Shabnam Hakimi, Jacob M. Parelman, Lun Yin, Jonathan R. Law, Jesse A.G. Skene, David A. Ball, Artemis Malekpour, Donald H. Beskind, Neil Vidmar, John M. Pearson, R. McKell Carter, J. H. Pate Skene

## Abstract

Efforts to explain complex human decisions have focused on competing theories emphasizing utility and narrative mechanisms. These are difficult to distinguish using behavior alone. Both narrative and utility theories have been proposed to explain juror decisions, which are among the most consequential complex decisions made in a modern society. Here, we asked jury-eligible male and female subjects to rate the strength of a series of criminal cases while recording the resulting patterns of brain activation. We compared patterns of brain activation associated with evidence accumulation to patterns of brain activation derived from a large neuroimaging database to look for signatures of the cognitive processes associated with different models of juror decision making. Evidence accumulation correlated with multiple narrative processes, including reading and recall. Of the cognitive processes traditionally viewed as components of utility, activation patterns associated with uncertainty, but not value, were more active with stronger evidence. Independent of utility and narrative, activations linked to reasoning and relational logic also correlated with increasing evidence. Hierarchical modeling of cognitive processes associated with evidence accumulation supported a more prominent role for narrative in weighing evidence in complex decisions. However, utility processes were also associated with evidence accumulation. These complementary findings support an emerging view that integrates utility and narrative processes in complex decisions.

**Significance Statement:** The last decade has seen a sharply increased interest in narrative as a central cognitive process in human decision making and as an important factor in the evolution of human societies. However, the roles of narrative versus utility models of decision making remain hotly debated. While available models frequently produce similar behavioral predictions, they rely on different cognitive processes and so their roles can be separated using the right neural tests. Here, we use brain imaging during mock juror decisions to show that cognitive processes associated with narrative, and to a lesser extent utility, were engaged while subjects evaluated evidence. These results are consistent with interactions between narrative and utility processes during complex decision making.

## Introduction

Narrative models have gained traction as an alternative to traditional utility models in economics (Shiller, 2017), psychology (Tuckett and Nikolic, 2017), and evolutionary biology (Gottschall et al., 2005; Wiessner, 2014) as a way of understanding real-world human decisions (Cialdini and Goldstein, 2004; Thaler and Sunstein, 2009; Shiller, 2017; Tremblay et al., 2017; Menon, 2018). While both narrative and utility models describe a process in which people evaluate evidence to reach a decision, the two models have traditionally invoked different cognitive processes to explain evidence evaluation. Historically in economics, utility is more strongly associated with quantitative processing (von Neumann and Morgenstern, 1944), while narrative is associated with the goodness of fit of the current details to memories and mental schema (Shiller, 2017). As these models have evolved, they have begun to overlap conceptually. To test whether these models are separable or need to be synthesized, we seek to identify the cognitive processes active during evidence accumulation.

The (similarities and) differences between narrative and utility models have been explored extensively in the context of juror decisions in the criminal justice system (Arkes and Garske, 1982; Devine, 2012; Allen and Pardo, 2019; Hastie, 2019). In the context of criminal trials, utility theory assumes that jurors assess the cumulative weight of the evidence to estimate the probability that the defendant committed the crime, and compare that probability to a threshold based on expected harms or benefits (Kaplan, 1968; Connolly, 1987; Arkes et al., 2012; Devine, 2012). Thus, the cognitive processes most central to utility models are calculations of the likelihood or uncertainty of different events and the value of alternative outcomes (Camerer et al., 2005; Glimcher and Fehr, 2013; Dennison et al., 2022). For jury decisions, narrative explanations are based on the extent to which the total body of information about a case can be structured into a coherent account and how closely the competing narratives offered by prosecution and defense match the juror’s background knowledge, experience, and beliefs (Arkes and Garske, 1982; Pennington and Hastie, 1986; Devine, 2012; Allen and Pardo, 2019; Hastie, 2019). Narrative models traditionally emphasize cognitive processes that support language, semantic representations, and episodic memory (AbdulSabur et al., 2014; Britton and Pellegrini, 2014; Baldassano et al., 2018).

Here, we use neural data to test for the involvement of cognitive processes that historically distinguish utility or narrative models of evidence processing. For this, we use a previously characterized task in which participants evaluate the strength of the prosecution’s evidence against a criminal defendant (Pearson et al., 2018). By relying on non-overlapping sets of cognitive processes to characterize the two models (and not any common processes like early visual cognition) we preserve the ability to weigh evidence in favor of each model. We recognize, however, that in modern conceptualizations, some cognitive processes are shared between utility and narrative frameworks. We therefore use a hierarchical family of models to allow us to identify individual cognitive processes that may form the basis of a synthesis of the two models. For completeness, we also include social and affective processes associated with juror decisions.

Using whole brain decoding (Li et al., 2017), we compare patterns of brain activation associated with those cognitive processes to activation patterns observed with parametric evidence accumulation in a criminal case. We extend the whole-brain decoding approach by comparing the goodness of fit for each model of juror decision making. This new approach allows us to begin to resolve the contributions of each psychological model to a given decision.

## Materials and Methods

### Subjects, recruitment, and sampling

This manuscript analyzes different models of evidence accumulation for a dataset previously reported in (Castrellon et al., 2021); the previous study examined models of evidence-independent bias in juror decision making. Thirty-three healthy jury-eligible adults between the ages of 18 and 52 were recruited from the Durham, NC community and underwent a single-session fMRI scan. All participants were screened for significant health or neurological problems and had normal or corrected-to-normal vision, and all gave written informed consent before the start of the experiment. Neuroimaging data from four participants were excluded from the analysis because of technical issues. Twenty-nine participants (mean age = 30.6 ± 9.58 years, 15 women and 14 men) remained for neuroimaging analyses. The experiment was approved by the local ethics committee at Duke University Medical Center.

### Experimental procedures

The mock-juror task (Fig. 1A), answering a call to use engaging narratives in neuroscience (Willems et al., 2020), was adapted from earlier work (Pearson et al., 2018) for use in the fMRI scanner. The task asks participants to evaluate the strength of the case presented by the prosecution against a criminal defendant, representing the first phase of a criminal trial. We did not include any evidence that might be part of the defense case. Our task therefore reflects the accumulation of evidence against the accused, but does not address processes involved in evaluating conflicting evidence.

**Fig. 1.**
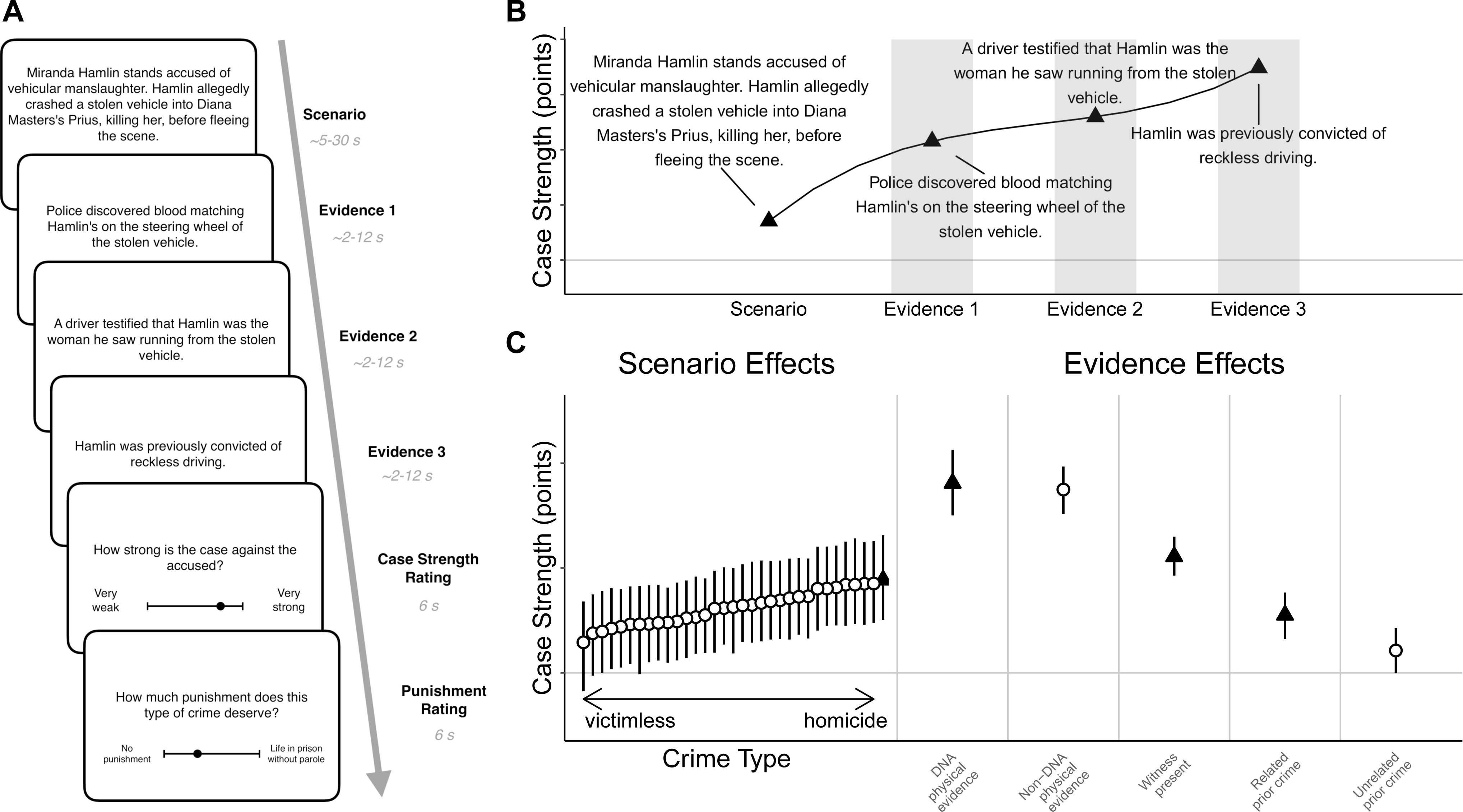
A mock-juror task separates evidence accumulation and crime-type bias. During the task, participants view a series of screens in the order depicted (**A**), first describing a criminal scenario followed by variable evidence from each of three categories (Extended Data Figure 1-1); following the evidence presentations, participants rate the strength of the case and the recommended punishment severity. (**B**) An example combination of scenario and evidence types is plotted as individual filled triangles for the accumulation of mean case-strength ratings (scale 0-100). Shaded regions indicate events included in the parametric regressor modeling the accumulation of evidence. (**C**) Case-strength contributions from scenario independent of evidence and case-strength contributions from evidence independent of scenario. Symbols represent mean effect size (scale 0-100); error bars represent 95% credible intervals. Scenario and evidence types depicted in panel B are distinguished with triangles. Scenario effects are shown in rank order.

Because of the large number of combinations of scenarios and evidence, these effects must be estimated across subjects. Subjects therefore viewed 33 scenarios, each describing an accusation of a specific crime that span shoplifting to rape, murder and child sexual abuse. Each crime scenario was followed by up to 3 items of evidence drawn at random from the evidence categories and levels described below. The task was split into three runs with 10-12 trials (scenarios) per run. For each scenario, participants first viewed a textual description of a crime type (a ‘scenario’) presented for 5-30s. This initial description contains no evidence of who committed the crime. While type of crime is the primary difference between scenarios, the scenarios also differ in other details, including names of defendants and victims, and the circumstances of the crime. Varying these other details is designed to keep subjects engaged throughout the task and to encourage participants to treat each scenario as a distinct crime. At the same time, the varied descriptions raise the potential for differences other than crime type to influence effects of the crime scenario on judgments of guilt. However, our large number of scenarios and the range of crime seriousness across those scenarios allows us to test specifically for a main effect of crime seriousness on judgments of guilt. Scenario and evidence texts are from (Pearson et al., 2018) and are available in Extended Data (Fig 1-1).

After viewing the crime scenario description, participants are shown three types of evidence implicating the named suspect (text presented one sentence at a time across multiple screens for 2-12s). Subjects viewed evidence that was: (1) physical evidence, (2) eyewitness, and (3) criminal history. The order of presentation for these evidence types were randomized. Evidence possibilities that link the accused to the crime included (1) one of three options of physical evidence (no physical evidence, DNA evidence or non-DNA physical evidence such as fingerprint or ballistic evidence); (2) either of two levels of eyewitness (eyewitness or a non-eyewitness), and (3) one of three options of criminal history (no prior convictions, prior conviction for a related crime, or prior conviction for an unrelated crime). Each participant saw one randomized combination of evidence with each of 33 crime scenarios. See Figure 1-1 for all possible variations. Following presentation of the scenario and evidence, participants were asked to rate the strength of the case and recommend a degree of punishment (on a scale of 0-100) for 6s each. Across subjects, there are 18 unique evidence combinations (3 × 2 × 3) for each crime scenario. Subjects may not view each of the 18 combinations of evidence but the effects of each individual piece of evidence can be estimated across subjects and across scenarios.

### Computational modeling

We used a hierarchical Bayesian model to estimate scenario-level and evidence-level effects on case strength and punishment ratings. Since this model has the advantage of accounting for sparsely sampled data, we can estimate effects for all scenarios even though participants did not view all possible combinations of scenarios and evidence types. We previously applied this model to a large online sample of participants (Pearson et al., 2018) and show here that the model generally replicates case strength and punishment effects (Castrellon et al., 2021).

To account for both case strength and punishment ratings for a given scenario, we model the resulting vector of ratings, *R_r_, r* = 1…*N_r_*, similarly:

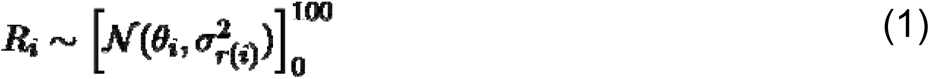

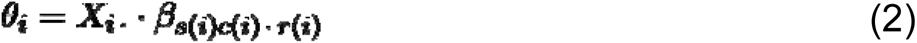

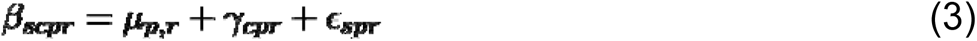

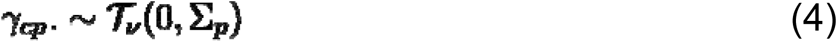

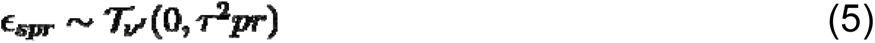

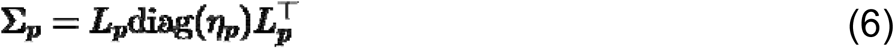

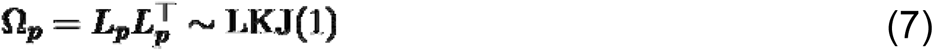

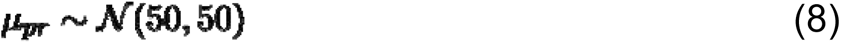

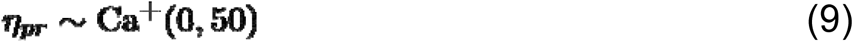

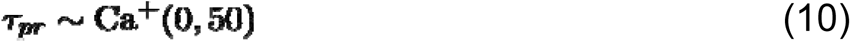

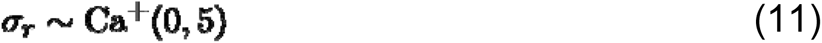

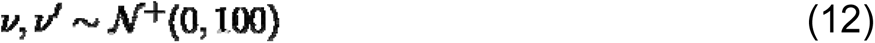

Here, we have used a "long" or "melted" representation of in which each index *i* corresponds to a single observation of a single rating scale *r(i)*. This allows us to more easily handle missing data in the model fitting procedure (see below). Here, (4) and (7) involve a multivariate t-distribution on the population effects specific to each case (*γ*) That is, we allow for covariance among the ratings for each effect at the population level, where the magnitudes of the variances are again controlled by *η* and the correlations Ω = *LL*^T^ are modeled according to the LKJ distribution (Lewandowski et al., 2009) through their Cholesky factorization (7). Second, in order to more accurately estimate variances in the presence of missing data, we have restricted this model to a single value of Τ across all cases (for each outcome and regressor) (3).

We calculated posterior distributions and credible intervals for each variable of interest using Markov Chain Monte Carlo methods as implemented in the Stan probabilistic programming language (Carpenter et al., 2017). Full code for all analyses is available at https://github.com/jcastrel/juror_fmri. Models were fitted by running 4 chains of 2000 samples with a thinning fraction of 2. The first half of samples were discarded as burn-in. This resulted in 2000 total samples for each variable, for which we report means as well as 95% equal-tailed credible intervals (bounded by the 2.5% and 97.5% samples from the distribution). We assessed convergence via effective sample size and the Gelman-Rubin statistic, for which all runs had 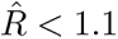 (Gelman et al., 2014).

Since the model weights for crime-type bias (case strength and punishment for scenario) across the 33 scenarios were highly correlated in this sample (r = 0.904, p < 0.001) inclusion of both weights as parametric fMRI regressors would weaken our ability to separate both effects during scenario viewing. To compensate for this challenge, we ran a principal components analysis (PCA) to derive a common metric that explained sufficient shared variance of case strength and punishment (PC1) and orthogonal residual shared variance (PC2). Therefore, PC1 represents a general measure of crime-type bias. In a separate dataset, we previously reported that crimes rated as deserving higher punishment were rated as having a higher case strength (regardless of evidence, (Pearson et al., 2018)). In that sample of Amazon mTurk participants (N = 360), case strength and punishment weights were also correlated (r = 0.389, p = 0.025). Prior to PCA, case strength and punishment weights were standardized (z-scored). In the fMRI sample, PC1 explained 95.2% of the variability and PC2 explained 4.80% of the variability. Similarly, in the mTurk sample, PC1 explained 69.5% of the variability and PC2 explained 30.5% of the variability.

### fMRI data acquisition

Brain images were collected using a 3T GE MR750 with an 8-channel head coil. For each of the 3 runs of the mock juror task, we acquired a T2*-weighted spiral-in sensitivity-encoding sequence (acceleration factor = 2) with slices parallel to the axial plane connecting the anterior and posterior commissures to acquire 488 volumes of 37 ascending interleaved slices, in-plane resolution = 3.75 x 3.75 x 3.75 mm, acquisition matrix = 64 x 64 and FOV = 243 x 243 mm, flip angle = 60°, TR = 1580 ms, TE = 30 ms. Before preprocessing these functional data, we discarded the first 7 volumes of each run to allow for stabilization of the T2* signal. To facilitate coregistration and normalization of these functional data, we also acquired whole-brain high-resolution anatomical scans (T1-weighted fast spoiled gradient echo sequence, in-plane resolution 1 × 1 × 1 mm; acquisition matrix = 256 × 256, FOV = 256 x 256 mm; 206 axial slices; flip angle = 12°, TR, 8.16 ms; TE, 3.18 ms).

### fMRI data preprocessing

Preprocessing was performed using FSL software (version 5.0.1). Functional data were motion-corrected with MCFLIRT and aligned to the middle volume in the time series (Jenkinson et al., 2002). Non-brain tissue from anatomical and functional data was removed using BET (Smith, 2002). Functional data were high-pass filtered with a cutoff of 100 seconds, spatially-smoothed with a 6 mm FWHM Gaussian kernel, and grand-mean intensity normalized. Functional images from each run were coregistered to subject’s anatomical images using FLIRT with 6 degrees of freedom and nonlinearly transformed to a standard MNI space 2-mm^3^ template using FNIRT with 12 degrees of freedom (Jenkinson and Smith, 2001).

### Univariate fMRI analysis

Statistical modeling of fMRI data was performed using FSL FEAT (version 6.0.0) (Smith et al., 2004; Woolrich et al., 2009). First-level analyses used FILM prewhitening for autocorrelation correction. Event variables and parametric regressors were convolved with a double-gamma hemodynamic response function. For each subject, a general linear model was fit to the data with mean event regressors for 1.) scenario viewing, 2.) evidence viewing, 3.) case strength rating, and 4.) punishment rating. To identify regions associated with modulation of BOLD signal with crime-type bias, we included two parametric regressors for loading scores on the principal components (crime-type bias/PC1 and PC2) from the PCA described above between case strength and punishment model weights.

Evidence accumulation was modeled as a parametric regressor of the cumulative sum of evidence weights as each piece of evidence was presented. For each trial, the evidence accumulation model starts at zero and then is incremented when a piece of evidence is presented. The regressor is incremented at the onset of presentation of each piece of evidence by an amount corresponding to the mean population weight from the behavioral analysis. The regressor is reset to zero at the end of the evidence presentation period for each trial. In Fig. 1B, for example, the regressor is zero after the scenario presentation and increments to 36.18 after the presentation of the first item of evidence (DNA), to 58.36 (plus 22.18) after the second evidence item (eyewitness), and to 69.39 (plus 11.03) after the third item of evidence (prior conviction). It then resets to zero at the as the third item is removed from the screen.

Additional parametric regressors included the scenario-level case strength model weights during the case strength rating event and scenario-level punishment model weights during the punishment rating event. Second-level analyses averaged within-subjects’ data across runs using a fixed-effects model. Higher-level analysis used a mixed-effects model (FLAME1) to average data across subjects. Statistical maps were thresholded using a cluster-forming threshold with a height of Z > 2.3, and cluster-corrected significance threshold of p < 0.05.

### Neurosynth Similarity Analysis

To evaluate the quality of our topic models in an unbiased and data-driven fashion, we followed up the linear regression with topic-based spatial correlation analysis. Here, we estimated the spatial voxelwise Pearson correlation between our parametric fMRI maps and each of all the possible 200 topic maps in the Neurosynth database. We visualized these correlations using a histogram for the thresholded map (Fig. 3). We expected the specific topics with the strongest associations would match those from our a priori models.

**Fig. 3.**
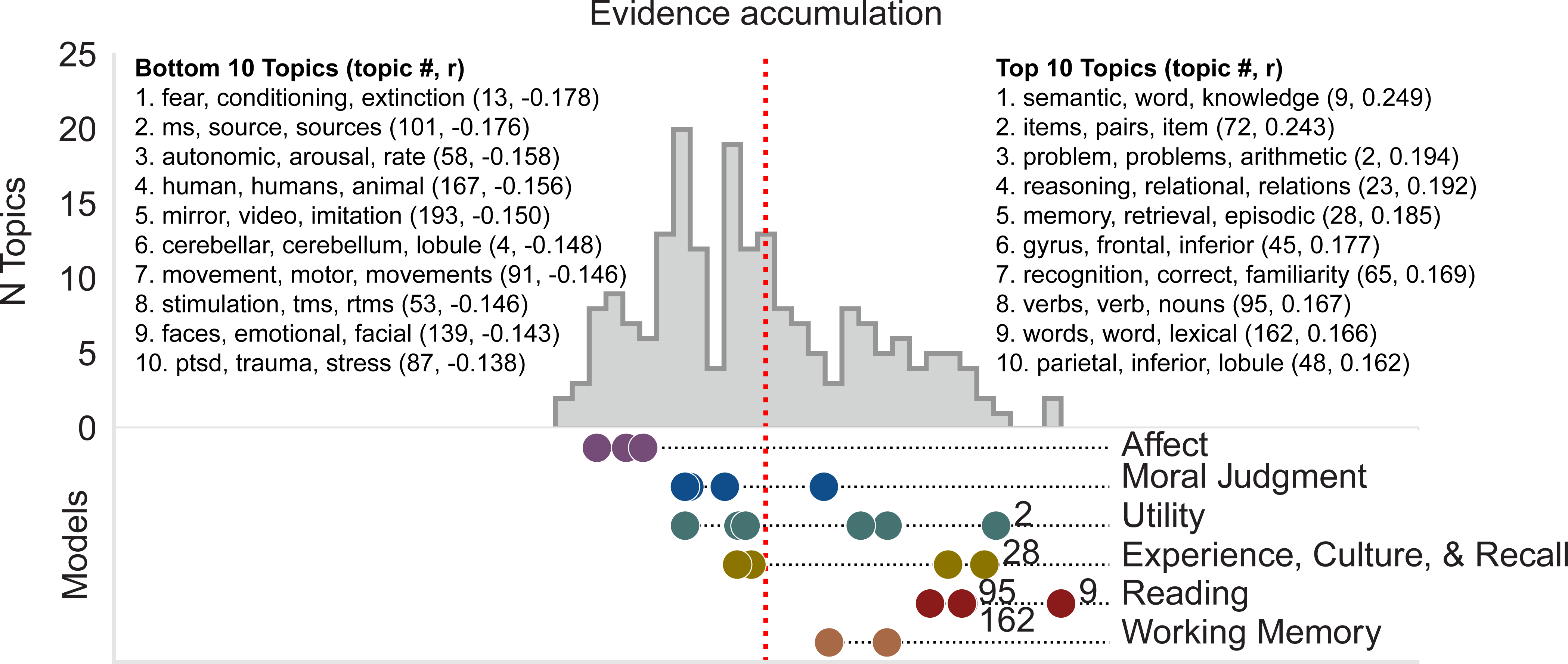
Brain activation patterns associated with evidence accumulation are associated with different cognitive processes. Histogram represents the distribution of spatial correlations between 200 Neurosynth topic maps and fMRI maps for evidence accumulation. Below the histogram are points that correspond to the location of the correlation coefficients in the histogram for each topic included in each of the indicated decision-making models. The top and bottom 10 correlated topics from the entire set of 200 topics are indicated beside the histograms (top panels) and topics within each model that appear in the top 10 list are labeled beside their associated point for each model (bottom panel). Within the reading model, the dots for topics 95 and 162 overlap.

To facilitate comparison of utility and narrative frameworks, we grouped cognitive topics from Neurosynth into a hierarchical set of cognitive models and submodels (preregistered at https://osf.io/rk92x/). To preserve the ability to compare models, we constructed the groups of cognitive processes as non-overlapping sets. At the most general level, we distinguish between evidence-based and social-affective based models of decision making. The evidence group is then separated into the utility and narrative frameworks we compare here. Within the utility model, we included topics associated with the numeric estimation of value and uncertainty. Within the narrative framework, we included topics associated with memory recall, reading comprehension, and working memory. Within the social-affective framework, we included topics that do not logically inform the decision of case-strength, including social cognition, affective reaction, and moral judgment. For each model and submodel, we first identified Neurosynth topics that contained terms related to the target cognitive processes; we then reviewed the studies included in each Neurosynth topic to confirm that they are consistent with the topic. The list of topics included in each model is provided in Figure 4-1.

**Fig. 4.**
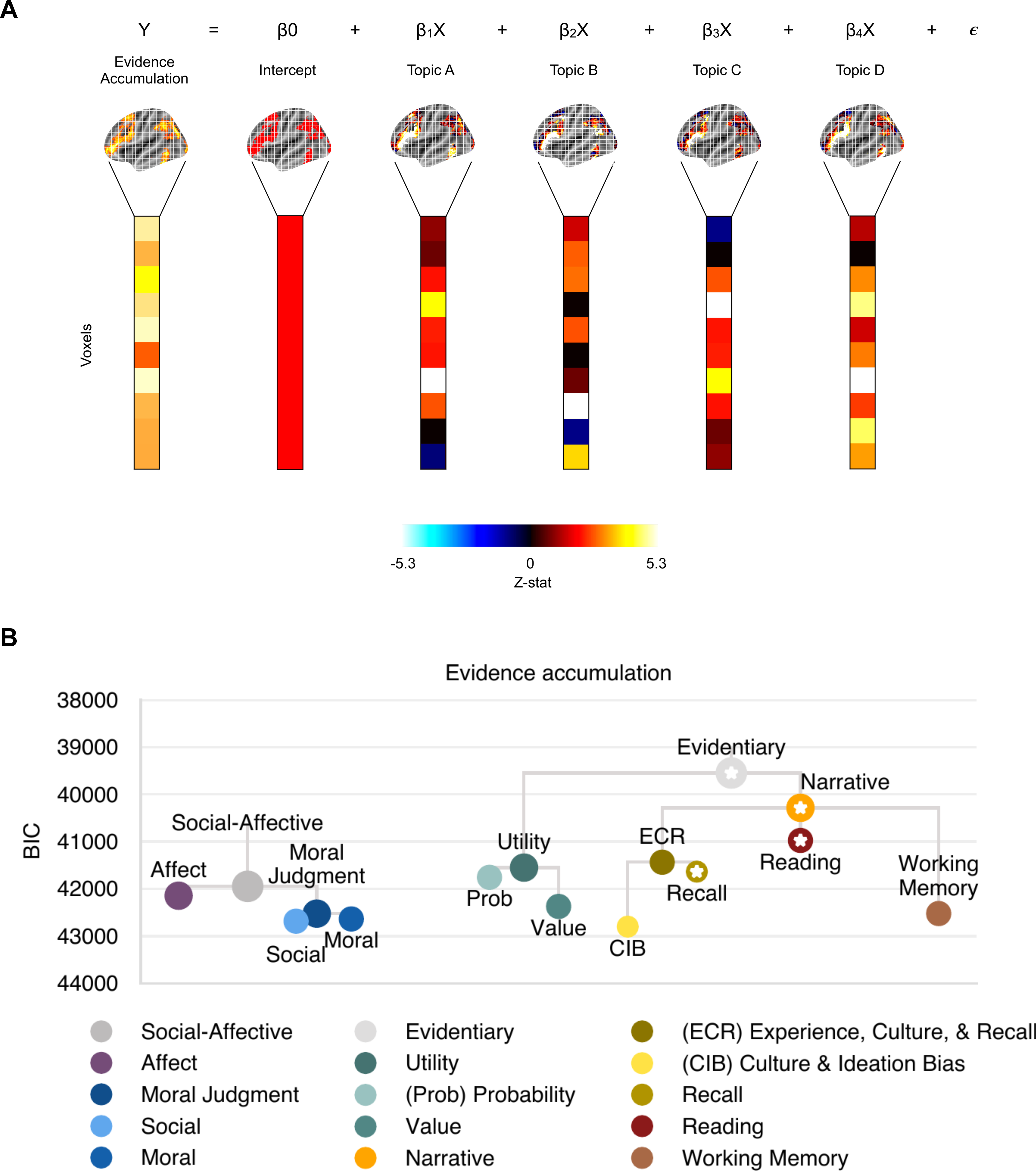
Patterns of brain activation associated with evidence accumulation support narrative-based processing. **(A)** Illustration of model comparison using spatial similarity analysis. To determine which proposed models of juror decision making best explain evidence accumulation, we performed a spatial voxelwise linear regression between evidence accumulation and Neurosynth topic maps associated with a given decision making model. We constructed an inclusion mask of voxels that were significantly positively associated with evidence accumulation. We then used that map to select voxels from the Neurosynth maps and the parametric map for evidence accumulation. Voxel values were vectorized into one-dimensional arrays and entered into a linear regression. The colorbar represents positive (warmer colors) or negative (cooler colors) statistical values. Neurosynth topic-map values could be positive or negative. **(B)** Height of dendrograms shows relative differences in fit (BIC) of Neurosynth topic models and submodels (topic list in Extended Data Figure 4-1) to the parametric fMRI evidence accumulation map (parameter estimates in Extended Data Figure 4-2). Lower BIC scores indicate better model fit. Stars indicate models with BIC scores that are lower than the 95 percentile of equally-sized random models (thresholds in Extended Data Figure 4-3).

Specifically, we grouped meta-analytic reverse-inference fMRI statistical maps from the Neurosynth database (version 5, 200 topics release: https://neurosynth.org/analyses/topics/v5-topics-200/) (Yarkoni et al., 2011) that reflect cognitive features associated with distinctive models of juror decision making. For example, a moral judgment model included Neurosynth maps from topic 112 (“context”, “empathy”, “prosocial”, etc.), topic 135 (“moral”, “guilt”, “harm”, etc.), topic 145 (“mind”, “mental”, “social”, etc.), and topic 154 (“social”, “interactions”, “interpersonal”, etc.). We grouped maps according to a hierarchical structure with primary models and nested submodels. For the moral judgment model, for example, topic maps 112 and 135 contributed to a moral submodel and topic maps 145 and 154 contributed to a social submodel. For the utility model, we included core cognitive topics associated with value, the consideration of risk, and mathematical calculation. For the narrative model, we included a series of submodels: 1. Experience, culture, and recall narrative (meant to encompass models associated with the consistency of a story and past experiences). This model was further subdivided into: 1a. culture and ideation bias and 1b. recall; 2. Working memory narrative (meant to capture the process of holding the pieces of a story together); and 3. Reading narrative (meant to capture models associated with processing fluency). For social-affective models, we considered both affect and moral judgment. Within moral judgment, we included submodels for the process of applying morality and, separately, broader studies of social cognition.

After defining the models, we ran spatial linear regression using the topic maps as predictors of each model’s similarity to our parametric statistical maps for evidence accumulation using the form 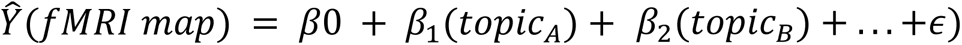 for *i* topics included in each model. We used unthresholded z-statistic maps from our results and unthresholded z-statistic Neurosynth topic maps provided by the Neurosynth developer Tal Yarkoni (since the viewable maps on the Neurosynth website are thresholded). We restricted inclusion to voxels that survived cluster-based thresholding for positive parametric effects (Fig. 2). While negative activations could potentially play an important role in juror decision making, it is also possible that negative activations reflect merely a minimization of energy use by suppressing task-irrelevant processes. Future analysis of negative activations will require a task that includes both inculpatory and exculpatory evidence.

**Fig. 2.**
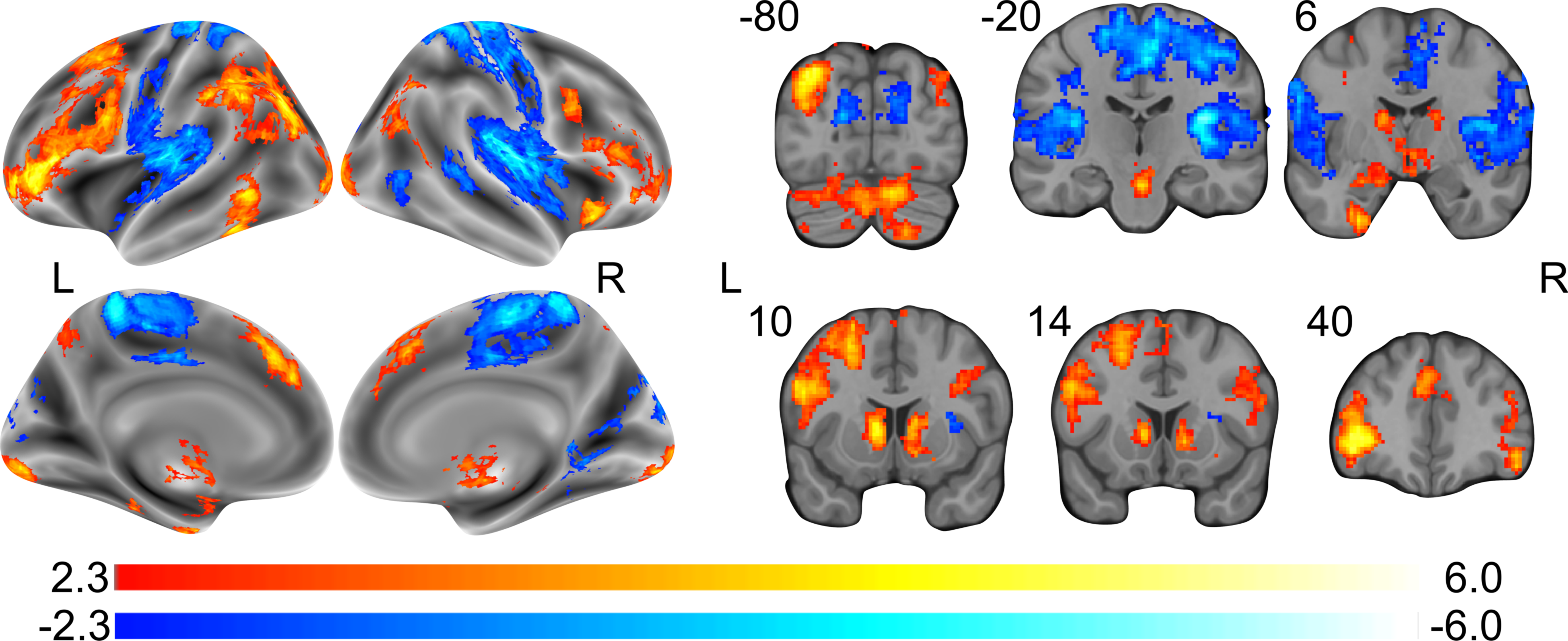
Evidence accumulation modulates neural activity patterns. Univariate fMRI patterns are shown for the effect of evidence accumulation during evidence presentation. Results shown are corrected for multiple comparisons using a whole brain cluster-forming threshold Z > 2.3, cluster-corrected *p* < 0.05.

FMRI spatial maps were vectorized into one-dimensional arrays using the ‘nibabel’ package. We ran linear regressions using the ‘OLS’ and ‘fit’ functions from the ‘statsmodels’ Python package and extracted Bayesian information criterion (BIC) scores. Lower BIC scores indicate greater similarity of the fMRI spatial maps to the decision-making models.

To identify meaningful associations, we estimated a model-specific lower bound threshold of BIC scores by comparing each decision making model BIC score to a null model that consisted of topic maps that were not included in any a priori model (177/200 remaining topic maps). The number of topic maps included in a null model matched the number of topic maps in a given model. We resampled 5,000 combinations of the randomly selected topic maps without replacement. Our lower bound threshold was defined as the 95 percentile of null model BIC scores (indicating a model fit was higher than 4,750 null models (p <.05). Model fit and regression coefficients are presented in Figure 4-2.

We note that this analysis is similar to testing for representational similarity of topic features with activation in those areas that respond to accumulating evidence.

### Analysis of activation associated with reading

The model-based analysis suggested that a reading model explained a significant amount of the variability in neural activation associated with evidence accumulation. We considered the possibility that the association with reading reflects processing of textual features, such as word count or reading complexity, as opposed to the meaning and coherence of the narrative interpretation of the text.

Using FSL FEAT, we performed a least-squares analysis (Jenkinson et al., 2002; Smith, 2002) on the preprocessed fMRI data such that each participant’s data was modeled with a design matrix composed of 132 columns for each trial’s event period with a single event in each column (33 scenarios x 4 event types: scenario, evidence (which includes all 3 evidence presentations), case strength rating, and punishment rating). These single-trial estimates were modeled with a double-gamma hemodynamic response function and the contrast representing the mean effect was extracted as a Z-statistic image. We then estimated the mean activation associated with the evidence reading events for each trial from a mask of the significant clusters associated with evidence accumulation (Fig 2).

To rule out the possibility that neural activation associated with evidence accumulation is driven by low-level textual features, we applied a mixed-effects model with a random intercept for each participant using the “lme4” and “lmerTest” packages in R to test the trial-level effect of summed word count from evidence pieces on fMRI activation. Using the “quanteda” R package, we calculated the number of words, syllables, and sentences for the evidence portion of each trial. We estimated reading complexity using the Flesch-Kincaid formula (F-K = (0.39*words per sentence) + (11.8*syllables per word) - 15.59) (Kincaid et al., 1975).

### Data and code availability

Unthresholded fMRI statistical maps can be viewed/downloaded from Neurovault: https://neurovault.org/collections/UADNVKNI/. Code and fMRI analysis preregistration for hypothesis grouping can be viewed and downloaded from OSF: https://osf.io/rk92x/ and Github: https://github.com/jcastrel/juror_fmri.

## Results

Jury-eligible participants viewed a series of simplified crime scenarios that vary in the type of crime and amount of evidence supporting guilt (Pearson et al., 2018) during functional magnetic resonance imaging (fMRI). Following each scenario, participants rated the strength of the case against the defendant and the punishment deserved for the crime. We used a hierarchical Bayesian model to estimate the effects of each type of evidence on participant ratings of case strength and punishment, controlling for the effects of different crime scenarios. Overall strength of the case reflected the sum of the weights of the individual pieces of evidence presented (Fig. 1). Here, we examine cognitive processes associated with evidence accumulation. We address the effects of different crime scenarios in a separate manuscript (Castrellon et al., 2021).

### Neural activations associated with evidence accumulation

We first sought to identify the patterns of brain activation associated with evidence accumulation. We modeled evidence accumulation as a parametric regressor of the cumulative sum of evidence weights as each piece of evidence was presented. We observed a constellation of activations associated with evidence accumulation, including distinct regional activations that were positively and negatively correlated with the cumulative weight of the evidence presented for each scenario (Fig. 2). Increasing strength of evidence evoked activation in the posterior temporal-parietal junction, dorsal-lateral prefrontal cortex, dorsal medial prefrontal cortex, ventral striatum, middle caudate, entorhinal cortex, hippocampus and amygdala. Decreasing strength of evidence was associated with activation in somatosensory, motor, early visual, and insular cortex as well as superior temporal sulcus. The unthresholded statistical map for evidence accumulation is accessible on Neurovault (https://neurovault.org/collections/UADNVKNI/).

The brain regions identified in this analysis have been linked to many different cognitive processes. To clarify the cognitive processes linked to evidence accumulation, we employed a data-driven approach, comparing the pattern of brain activations associated with evidence accumulation to the activations associated with each of 200 topics defined in the Neurosynth database (version 5, 200 topics release: https://neurosynth.org/analyses/topics/v5-topics-200/) (Yarkoni et al., 2011). Neurosynth topic maps reflect meta-analyses of more than 14,000 fMRI studies (Yarkoni et al., 2011). We used a spatial voxelwise linear regression to compare activations associated with evidence accumulation to the map of activations associated with each Neurosynth topic (Fig. 3). For this analysis, we considered only brain regions positively correlated with evidence accumulation because our current task varies only the amount of affirmative evidence of guilt. Analysis of negative correlations would require a task that included exonerating evidence, in order to distinguish a role for negative correlations in evidence processing from energy minimization in the brain. Topics with the strongest correlations for evidence accumulation included language comprehension topics (Topic 9 - semantic, word, knowledge; Topic 95 - verbs, nouns, syntactic; Topic 162 - words, lexical, meaning), memory topics (Topic 72 - pairs, associative, accuracy; Topic 28 - memory, retrieval, episodic; Topic 65 - recognition, correct, familiarity), a quantitative reasoning topic (Topic 2 - problems, mathematics, calculation), and a reasoning topic (Topic 23 - reasoning, relational, logic). These topics appear to align with aspects of both utility and narrative models. To formally compare which models are supported by these topics, we preregistered a hierarchical model with groups of topics relevant for separate psychological models.

### Neural support for two aspects of a narrative framework

We preregistered a list of cognitive topics associated with utility, narrative, and social-affective models and submodels (https://osf.io/rk92x/) to limit user degrees of freedom while defining model structure. To compare these models, we used a spatial voxelwise linear regression to compare activations associated with evidence accumulation to the map of activations associated with each Neurosynth topic (Fig. 4A). For the utility model, we included core cognitive topics associated with value, the consideration of risk, and mathematical calculation. For the narrative model, we included a series of submodels: 1. Experience, culture, and recall narrative (meant to encompass models associated with the consistency of a story and past experiences). This model was further subdivided into: 1a. culture and ideation bias and 1b. recall; 2. Working memory narrative (meant to capture the process of holding the pieces of a story together); and 3. Reading narrative (meant to capture models associated with processing fluency). We have separated these utility and narrative processes according to a split between quantitative and qualitative reasoning. However, we note that modern efforts have proposed the integration of features of recall into utility models of decision making (Shadlen and Shohamy, 2016; Zhang et al., 2021). To preserve the ability to compare these models, we maintain the sets of cognitive processes as two, non-overlapping, groups of cognitive processes. To consider the impact of factors outside of narrative and utility, we also tested a social-affective model, including affect and moral judgment. Within moral judgment, we included submodels for the process of applying morality and broader studies of social cognition. Hereafter, a model is described as significant if it explains more variance than 95% of models composed of randomly drawn topics from the 177 topics not associated with one of the a priori models. The null distribution for each test was defined for a number of topics matched to each model (e.g., 5 drawn topics if the model being evaluated contained 5 topics).

If evidence accumulation is largely a matter of quantitative reasoning regarding the likelihood that an event occurred, we would expect that patterns of brain activation associated with evidence accumulation would be similar to topic maps from the utility model. On the other hand, if evidence accumulation relies on non-quantitative processes, like the coherence and plausibility of a narrative, then we would expect brain activation for evidence accumulation to correspond to topic maps from the narrative models. For example, the recall of past experiences and events is more likely to be necessary when evaluating narrative plausibility.

Patterns of brain activation during evidence accumulation were significantly correlated with topic maps from the narrative model (BIC = 40285, p = 0.014, resampled null, Fig. 4B) but not utility (BIC = 41550, p = 0.24, resampled null, Fig. 4B) or social-affective (BIC = 41947, p = 0.72, resampled null, Fig. 4B) topic sets. Of the narrative submodels, both the narrative recall and narrative reading topic maps explained a significant percentage of the variance (Fig. 4B, BIC = 41436, p = 0.019, resampled null and BIC = 40975, p = 0.020, resampled null, respectively). The group of reading topics included cognitive processes associated with both the reading of individual words and sentences and the integration of words and sentences to interpret the meaning of the text. Three of the 4 preregistered topics in the reading model (see above) were significantly associated with activation during evidence accumulation. The fourth topic (topic 93 - language, comprehension, narrative) was not in the top 5% of the 200 topics tested (Fig. 3), but was closely grouped with the other three (unlabeled red dot in Fig. 3). While each of the four topic maps is independently correlated with evidence accumulation, topic 9 (knowledge, word, semantic) is the most strongly correlated (Fig. 3). In fact, when aggregated into an overall reading model, the contribution of topic map 9 is three times larger than the next largest contributing topic map (β=0.147 vs. β=-0.048, Figure 4-2). Topic map 9’s outsized contribution to the model suggests that the association between reading processes and evidence accumulation is due to more narrative-like processes (meaning associated) rather than lower level text features. In the next section, we conduct additional tests to further clarify the nature of the reading/evidence accumulation association.

### Low level reading features are not associated with evidence

In our preregistered hierarchical narrative and utility models, we assigned reading topics to the narrative model because higher-order aspects of reading are central to narrative construction and interpretation (AbdulSabur et al., 2014; Britton and Pellegrini, 2014; Baldassano et al., 2018). We recognize, however, that lower-level features of reading (word recognition, grammar, syntax) might also correlate with differences in evidence in a way that does not directly distinguish narrative from utility models. Since our models included both higher- and lower-order aspects of reading, we are able to distinguish these possibilities. To test whether high- or low-level features of reading contribute to the evaluation of evidence, we conducted two analyses, asking (1) Does evidence accumulation correlate with low-level text features? and (2) Does all text that increases perceived case strength evoke activation of the reading areas associated with evidence accumulation?

We first tested whether low-level language features of our evidence descriptions were correlated with the average activation in areas that responded to increasing evidence. In that analysis, the activation of brain regions associated with the reading submodel were not correlated with the cumulative number of words or complexity/readability in the evidence text, but were correlated parametrically with the cumulative weight of the evidence (Fig. 4B). Specifically, our post-hoc ROI analysis indicated that low-level textual features did not drive observed evidence accumulation patterns of activity in reading-processing regions. Neither word count (F(1,932.42) = 0.063, p = 0.802) nor reading complexity (F(1,932.89) = 0.322, p = 0.571) were associated with fMRI activation during evidence viewing (Fig. 5).

**Fig. 5.**
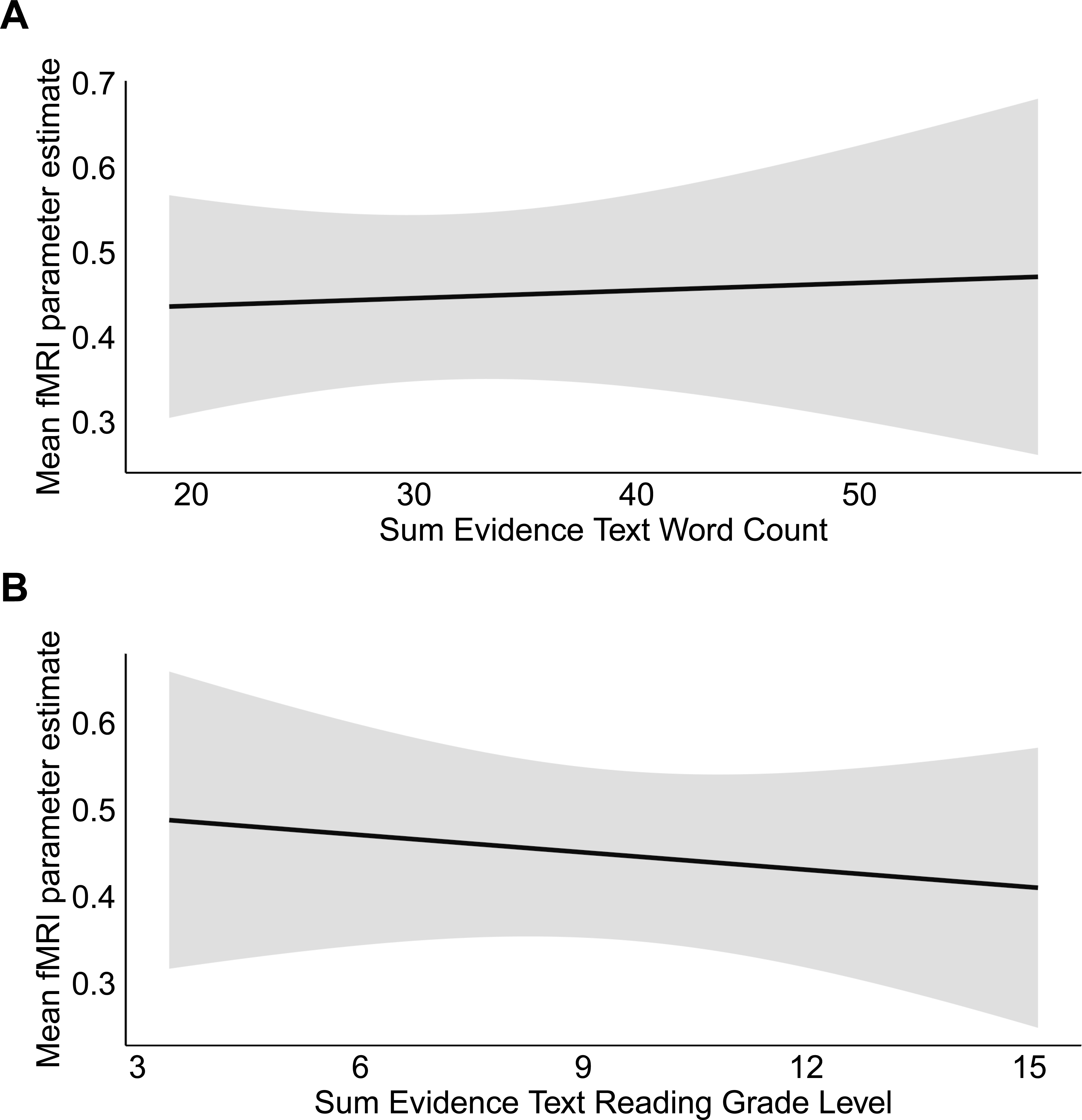
Low-level textual features do not account for fMRI activations that increase with stronger evidence. Trial-level estimated sum of (**A**) word count (F(1,932.42) = 0.063, p = 0.802) and (**B**) reading complexity (F(1,932.89) = 0.322, p = 0.571) of presented evidence text were not related to fMRI patterns associated with evidence accumulation. Shaded areas represent 95% confidence intervals.

This suggests that the activation of brain regions associated with reading is related to the interpretation rather than the length or complexity of the text. The correlation between patterns of brain activation observed with increasing evidence and patterns associated with recall provides support for the role of narrative in weighing evidentiary contributions to case strength.

### Reading activation is specific to evidence accumulation

To better understand the correlation of reading-topic maps with evidence accumulation, we tested whether activation of the reading topic maps is specific to evidence evaluation. Before reading the evidence in each case, participants first read a short scenario describing the crime (Fig. 1). We found previously that the severity of the crime described in each scenario affects case strength ratings independent of the evidence (Pearson et al., 2018). This allowed us to test whether scenario text, which also increases perceived case strength, evokes activation of the reading areas associated with evidence accumulation.

Elsewhere, we found that brain activation patterns associated with scenario effects on case strength are distinct from the patterns described here for evidence accumulation (Castrellon et al., 2021). These scenario effects were associated with brain regions related to social cognition. If the relationship between reading topics and evidence accumulation is a matter of simply reading any text that affects case strength, we would expect to find a significant relationship between the reading topic maps and increasing activation with scenario effects on case strength.

We therefore compared activation for scenario-driven case strength to the same reading topic maps used in our models of evidence accumulation. We found no significant relationship between increasing activation patterns associated with reading different crime scenarios and the reading topic maps (R^2^ = 0.013, p=0.646). We conclude that the association of reading topics with evidence accumulation is driven by higher-order features that distinguish evidence evaluation from other processes.

## Discussion

We have used neuroimaging to quantify the contributions of utility and narrative models to jury decision making, an example of complex human decisions. To do that, we extended a novel method for inferring cognitive processes from patterns of brain activation. Increasing levels of evidence evoked brain activation patterns associated with reading, recall, calculating uncertainty, and logic and reason. Reading and recall are typically associated with aspects of narrative, while uncertainty is traditionally associated with utility. In addition, the activation associated with logic and reason could be viewed as supporting either narrative or utility models, or as an entirely separate process. Nevertheless, our findings suggest a prominent role for the narrative model for evidence accumulation in juror decision making that could be extended to other complex decisions.

### Recall and reading processes track evidence accumulation

In addition to providing evidence of the importance of the narrative model, we were able to identify specific cognitive processes from the narrative model that significantly contributed to evidence accumulation in this task. We note that our analysis examined only the pattern of activity positively correlated evidence accumulation and does not address the negatively correlated regions (Fig. 2). Recall of past experiences (Baldassano et al., 2018) and reading are the only significant submodels that contribute to the explained variance. Recall allows the comparison of the current evidence to past experiences, stories, or schema (Baldassano et al., 2018), which can serve as a basis for assessing the plausibility and cumulative weight of the evidence.

Involvement of the reading submodel in evidence accumulation likely reflects the cohesion of the individual items of evidence rather than lower-level reading processes. The reading submodel includes not only the processes associated with reading individual words and sentences, but also the cognitive processes involved in integrating the content of words and sentences over time to assign meaning to the text. Consistent with this interpretation, imaging studies of subjects listening to stories have found that the integration of stories over increasing temporal windows shifts the pattern of brain activation from areas near auditory cortex to brain regions overlapping with those in our reading submodel (Lerner et al., 2011).

Previous work has shown that the familiarity and ease of processing a particular idea, including how easily it is read, is strongly associated with confidence in the validity of a statement (Hasher et al., 1977; Arkes et al., 1989; Fazio et al., 2015). Mentally, reading engagement might serve as an index of familiarity or the ease of understanding evidence that comprises a coherent account, and therefore confidence in the story presented, as proposed in the Story Model of juror decisions (Pennington and Hastie, 1986). Neuroimaging work comparing brain activation between true and false sentences generally supports this account, finding greater activation in the temporal parietal junction for true compared to false statements (Marques et al., 2009).

For this analysis, we focused on narrative as a mechanism for determining the likelihood that a particular story occurred (e.g., the story model (Pennington and Hastie, 1986)). As our models were formulated here, we find support for the narrative model in juror decision making. However, a significant body of research has studied narrative as an individual language process (Scholes, 1980; Britton and Pellegrini, 2014; Paulus et al., 2021), assembling information into memory schema (Baldassano et al., 2018; Chang et al., 2021), and communicating those schema to others (Shiller, 2017; Smith et al., 2017; Nguyen et al., 2020). Studies of narratives in juror decision making would benefit from connecting the aspects of evidence processing examined here to measures of narrative cohesion and how narrative schemas are encoded in the brain.

### Support for aspects of the utility model

We note that our findings do not exclude a role for quantitative reasoning in evidence accumulation. In fact, while the set of topics included in our utility model do not meet our statistical threshold, the model-independent analysis finds that arithmetic/quantitative terms are ranked third in terms of the variance in evidence accumulation explained (Fig. 3). This topic map overlaps with activation for evidence accumulation primarily in the intraparietal sulcus, an area associated with assessing uncertainty (Huettel et al., 2006; Levy et al., 2010). Topic maps associated with valuation overlap with activation for evidence accumulation only in the ventral striatum and not in the orbital frontal or ventral medial cortex (Hare et al., 2008; Bartra et al., 2013; Shadlen and Shohamy, 2016). Activation in ventral striatum is associated with both value and uncertainty (Knutson et al., 2005). Thus, ventral striatum is consistent with aspects of the utility (both value and uncertainty e.g. valuation (Bartra et al., 2013)) and narrative model (related to the probability of an event and prediction error (Abler et al., 2006; Rodriguez et al., 2006)). These findings support a role for uncertainty or probability estimation but do not support a role for value in evidence accumulation.

Support for aspects of utility as well as narrative processes in evidence accumulation raises the question of whether and how they could be integrated. Previous work has proposed to bridge utility and narrative frameworks by relying on processes associated with logic and reasoning (Johnson-Laird et al., 2015). Alternatively, memory recall could serve as a means of connecting narrative and utility. For example, work integrating memory and value posits a role for recall in utility calculations (Shadlen and Shohamy, 2016; Zhang et al., 2021).

In a narrative framework, recall is the proposed mechanism by which the likelihood of an event is estimated. Considering recall as a means of calculating likelihood would make it possible to reconfigure the utility model away from the conscious calculation of primarily numerical value to a more qualitative interpretation of utility. Such a shift might provide a means to merge aspects of the utility and narrative frameworks.

### Evidence and bias mechanisms are distinct

We previously found that the type of crime described in each of our scenarios influences participant ratings for the strength of the case, independent of the evidence (Pearson et al., 2018). Crime type and evidence accumulation thus represent distinct influences on case strength. Activation patterns associated with crime-type bias are located in a region of cortex associated with social cognition, including cultural and ideation bias (Castrellon et al., 2021). This crime-type activation is distinct from, but neighboring, those described here for the accumulation of evidence. The neighboring activations for recall and crime type bias are both located in a larger region hypothesized to combine information from different processing streams (Carter and Huettel, 2013; Jung et al., 2022). This is consistent with a narrative framework for integration of evidence, in which the accumulation of evidence is compared to prior experience and beliefs as a basis for decisions (Pennington and Hastie, 1986; Shiller, 2017; Hastie, 2019).

In this context, a further notable feature of evidence integration in the narrative model is the internal coherence of the evidence. In the current task, we investigated only cognitive processes associated with the accumulation of evidence consistent with the prosecution’s account of the crime. Our results therefore describe the process of evidence accumulation, but do not address whether the same or separate cognitive processes resolve conflicting information.

### Broader significance of narrative as a mechanism of decision making

Beyond the longstanding interest in narrative models in the context of juror decisions and trial practice (Pennington and Hastie, 1986; Lubet and Lore, 2015), narrative models have been offered as an alternative to standard utility models of decision making to help explain deviations from optimal behavior in a wide range of complex decisions from asset pricing to market bubbles (Shiller, 2017). Narrative models attempt to explain unusual behaviors like the nonlinear spread of preferences and the role of affect in decision making by assessing the cohesiveness of a story and comparing it to previous experiences. Work in anthropology and evolutionary biology suggest that storytelling has played an important role in human evolution as means of communicating plausibility based on past experience from generation to generation (Smith et al., 2017; Bietti et al., 2019; Hitchcock, 2019). In fact, recent work on the estimation of risk has found that human subjects are better at estimating risk when it is described as part of a story than when asked to do so numerically (Sinclair et al., 2021). It is therefore possible that in narratively structured settings, likelihood estimates can be constructed according to consistency with past experiences. This alternative means of assessing future (and past) likelihood represents a potentially significant expansion of how risk and uncertainty is treated in many complex decisions.

## Acknowledgements

The authors thank Tal Yarkoni for his assistance in accessing the Neurosynth database. This work was supported by an Incubator Award from the Duke Institute for Brain Sciences, The Institute of Cognitive Science at CU Boulder and a subcontract through eCortex, Inc. from the Office of Naval Research (ONR, N00014-18-C-2067) to R.M.C, National Science Foundation (NSF) grant no. 1655445 to J.H.P.S, and by a career development award from the NIH Big Data to Knowledge Program (grant no. K01-ES-025442 to J.M.P). J.J.C was supported by an NSF Graduate Research Fellowship (grant no. NSF DGE-1644868). The funders had no role in study design, data collection and analysis, decision to publish or preparation of the manuscript.

## Extended Data Legends

**Figure 1-1. Scenario crime descriptions and evidence options.**

**Figure 4-1. Neurosynth topic maps included in cognitive models and submodels.**

**Figure 4-2. Spatial similarity (linear regression fit) for juror decision-making models models and thresholded brain activation patterns associated with evidence accumulation and crime-type bias.** For evidence accumulation and crime-type bias fMRI maps, the table includes overall model fit (BIC score), p-value, and topic map standardized coefficient measures (beta and standard error (se)). P-values reflect the number of BIC values from the null distribution (n=5,000 permutations) that are greater than the model’s BIC.

**Figure 4-3. Null model spatial similarity threshold for juror decision-making models and thresholded brain activation patterns associated with evidence accumulation.** For evidence accumulation and crime-type bias fMRI maps, the table includes overall model fit (BIC score) threshold dependent on the number of topics included in a model. The threshold BIC is the 95 percentile of the null distribution of 5,000 random models.

## Bibliography

AbdulSabur, N. Y., Xu, Y., Liu, S., Chow, H. M., Baxter, M., Carson, J., et al. (2014). Neural correlates and network connectivity underlying narrative production and comprehension: a combined fMRI and PET study. Cortex 57, 107–127.

Abler, B., Walter, H., Erk, S., Kammerer, H., and Spitzer, M. (2006). Prediction error as a linear function of reward probability is coded in human nucleus accumbens. Neuroimage 31, 790–795.

Allen, R. J., and Pardo, M. S. (2019). Relative plausibility and its critics. The International Journal of Evidence & Proof 23, 5–59.

Arkes, H. R., and Garske, J. P. (1982). Psychological Theories of Motivation. Brooks/Cole.

Arkes, H. R., Hackett, C., and Boehm, L. (1989). The generality of the relation between familiarity and judged validity. J. Behav. Decis. Mak. 2, 81–94.

Arkes, H. R., Shoots-Reinhard, B., and Mayes, R. S. (2012). Disjunction Between Probability and Verdict in Juror Decision Making. J. Behav. Decis. Mak. 25, 276– 294.

Baldassano, C., Hasson, U., and Norman, K. A. (2018). Representation of Real-World Event Schemas during Narrative Perception. J. Neurosci. 38, 9689–9699.

Bartra, O., McGuire, J. T., and Kable, J. W. (2013). The valuation system: a coordinate-based meta-analysis of BOLD fMRI experiments examining neural correlates of subjective value. Neuroimage 76, 412–427.

Bietti, L. M., Tilston, O., and Bangerter, A. (2019). Storytelling as Adaptive Collective Sensemaking. Top. Cogn. Sci. 11, 710–732.

Britton, B. K., and Pellegrini, A. D. (2014). Narrative thought and narrative language. Psychology Press.

Camerer, C., Loewenstein, G., and Prelec, D. (2005). Neuroeconomics: How Neuroscience Can Inform Economics. J. Econ. Lit. 43, 9–64.

Carpenter, B., Gelman, A., Hoffman, M. D., Lee, D., Goodrich, B., Betancourt, M., et al. (2017). Stan: A probabilistic programming language. J. Stat. Softw. 76. doi: 10.18637/jss.v076.i01.

Carter, R. M., and Huettel, S. A. (2013). A nexus model of the temporal-parietal junction. Trends Cogn. Sci. 17, 328–336.

Castrellon, J. J., Hakimi, S., Parelman, J., Yin, L., Law, J. R., Skene, J. A., et al. (2021). Social cognitive processes explain bias in juror decisions. doi: 10.31234/osf.io/v56wz.

Chang, C. H. C., Lazaridi, C., Yeshurun, Y., Norman, K. A., and Hasson, U. (2021). Relating the Past with the Present: Information Integration and Segregation during Ongoing Narrative Processing. J. Cogn. Neurosci. 33, 1106–1128.

Cialdini, R. B., and Goldstein, N. J. (2004). Social influence: compliance and conformity. Annu. Rev. Psychol. 55, 591–621.

Connolly, T. (1987). Decision theory, reasonable doubt, and the utility of erroneous acquittals. Law Hum. Behav. 11, 101–112.

Dennison, J. B., Sazhin, D., and Smith, D. V. (2022). Decision neuroscience and neuroeconomics: Recent progress and ongoing challenges. *Wiley Interdiscip. Rev. Cogn. Sci.*, e1589.

Devine, D. J. (2012). Jury Decision Making: The State of the Science. NYU Press.

Fazio, L. K., Brashier, N. M., Payne, B. K., and Marsh, E. J. (2015). Knowledge does not protect against illusory truth. J. Exp. Psychol. Gen. 144, 993–1002.

Gelman, A., Carlin, J. B., Stern, H. S., Dunson, D. B., Vehtari, A., and Rubin, D. B. (2014). Bayesian Data Analysis (Chapman & Hall/CRC Texts in Statistical Science). Chapman and Hall/CRC.

Glimcher, P. W., and Fehr, E. (2013). Neuroeconomics: Decision Making and the Brain. Academic Press.

Gottschall, J., Wilson, D. S., and Wilson, E. O. (2005). The Literary Animal: Evolution and the Nature of Narrative. Northwestern University Press.

Hare, T. A., O’Doherty, J., Camerer, C. F., Schultz, W., and Rangel, A. (2008). Dissociating the role of the orbitofrontal cortex and the striatum in the computation of goal values and prediction errors. J. Neurosci. 28, 5623–5630.

Hasher, L., Goldstein, D., and Toppino, T. (1977). Frequency and the conference of referential validity. Journal of Verbal Learning and Verbal Behavior 16, 107–112.

Hastie, R. (2019). The case for relative plausibility theory: Promising, but insufficient. The International Journal of Evidence & Proof 23, 134–140.

Hitchcock, R. K. (2019). Hunters and gatherers past and present: Perspectives on diversity, teaching, and information transmission. Rev. Anthr. 48, 5–37.

Huettel, S. A., Stowe, C. J., Gordon, E. M., Warner, B. T., and Platt, M. L. (2006). Neural signatures of economic preferences for risk and ambiguity. Neuron 49, 765– 775.

Jenkinson, M., Bannister, P., Brady, M., and Smith, S. (2002). Improved optimization for the robust and accurate linear registration and motion correction of brain images. Neuroimage 17, 825–841.

Jenkinson, M., and Smith, S. (2001). A global optimisation method for robust affine registration of brain images. Med. Image Anal. 5, 143–156.

Johnson-Laird, P. N., Khemlani, S. S., and Goodwin, G. P. (2015). Logic, probability, and human reasoning. Trends Cogn. Sci. 19, 201–214.

Jung, H., Wager, T. D., and Carter, R. M. (2022). Novel Cognitive Functions Arise at the Convergence of Macroscale Gradients. J. Cogn. Neurosci. 34, 381–396.

Kaplan, J. (1968). Decision Theory and the Factfinding Process. Stanford Law Rev. 20, 1065–1092.

Kincaid, J. P., Fishburne, J., Robert P., R., Richard L., C., and Brad S. (1975). Derivation of new readability formulas (automated readability index, fog count and Flesch reading ease formula) for navy enlisted personnel. Fort Belvoir, VA: Defense Technical Information Center doi: 10.21236/ada006655.

Knutson, B., Taylor, J., Kaufman, M., Peterson, R., and Glover, G. (2005). Distributed neural representation of expected value. Journal of Neuroscience 25, 4806.

Lerner, Y., Honey, C. J., Silbert, L. J., and Hasson, U. (2011). Topographic Mapping of a Hierarchy of Temporal Receptive Windows Using a Narrated Story. Journal of Neuroscience 31, 2906–2915.

Levy, I., Snell, J., Nelson, A. J., Rustichini, A., and Glimcher, P. W. (2010). Neural representation of subjective value under risk and ambiguity. J. Neurophysiol. 103, 1036–1047.

Lewandowski, D., Kurowicka, D., and Joe, H. (2009). Generating random correlation matrices based on vines and extended onion method. J. Multivar. Anal. 100, 1989– 2001.

Li, R., Smith, D. V., Clithero, J. A., Venkatraman, V., Carter, R. M., and Huettel, S. A. (2017). Reason’s Enemy Is Not Emotion: Engagement of Cognitive Control Networks Explains Biases in Gain/Loss Framing. J. Neurosci. 37, 3588–3598.

Lubet, S., and Lore, J. C. (2015). Modern Trial Advocacy Analysis & Practice: Fifth Edition (NITA). 5 edition. Wolters Kluwer.

Marques, J. F., Canessa, N., and Cappa, S. (2009). Neural differences in the processing of true and false sentences: insights into the nature of “truth” in language comprehension. Cortex 45, 759–768.

Menon, A. (2018). Bringing cognition into strategic interactions: S trategic mental models and open questions. Strategic Manage. J. 39, 168–192.

Nguyen, M., Chang, A., Micciche, E., Meshulam, M., Nastase, S. A., and Hasson, U. (2020). Teacher-student neural coupling during teaching and learning. bioRxiv, 2020.05.07.082958. doi: 10.1101/2020.05.07.082958.

Paulus, I. D. P. C., Charest, I. D. I., and Benoit, I. D. R. G. (2021). Value shapes the structure of schematic representations in the medial prefrontal cortex. biorxiv. doi: 10.1101/2020.08.21.260950.

Pearson, J. M., Law, J. R., Skene, J. A. G., Beskind, D. H., Vidmar, N., Ball, D. A., et al. (2018). Modelling the effects of crime type and evidence on judgments about guilt. Nat Hum Behav 2, 856–866.

Pennington, N., and Hastie, R. (1986). Evidence evaluation in complex decision making. J. Pers. Soc. Psychol. 51, 242–258.

Rodriguez, P. F., Aron, A. R., and Poldrack, R. A. (2006). Ventral-striatal/nucleus-accumbens sensitivity to prediction errors during classification learning. Hum. Brain Mapp. 27, 306–313.

Scholes, R. (1980). Language, Narrative, and Anti-Narrative. Crit. Inq. 7, 204–212.

Shadlen, M. N., and Shohamy, D. (2016). Decision Making and Sequential Sampling from Memory. Neuron 90, 927–939.

Shiller, R. J. (2017). Narrative Economics. Am. Econ. Rev. 107, 967–1004.

Sinclair, A. H., Hakimi, S., Stanley, M. L., Adcock, R. A., and Samanez-Larkin, G. R. (2021). Pairing facts with imagined consequences improves pandemic-related risk perception. Proc. Natl. Acad. Sci. U. S. A. 118. doi: 10.1073/pnas.2100970118.

Smith, D., Schlaepfer, P., Major, K., Dyble, M., Page, A. E., Thompson, J., et al. (2017). Cooperation and the evolution of hunter-gatherer storytelling. Nat. Commun. 8, 1853.

Smith, S. M. (2002). Fast robust automated brain extraction. Hum. Brain Mapp. 17, 143–155.

Smith, S. M., Jenkinson, M., Woolrich, M. W., Beckmann, C. F., Behrens, T. E. J., Johansen-Berg, H., et al. (2004). Advances in functional and structural MR image analysis and implementation as FSL. Neuroimage 23 Suppl 1, S208–19.

Thaler, R. H., and Sunstein, C. R. (2009). Nudge: Improving Decisions about Health, Wealth, and Happiness. Penguin.

Tremblay, S., Sharika, K. M., and Platt, M. L. (2017). Social Decision-Making and the Brain: A Comparative Perspective. Trends Cogn. Sci. 21, 265–276.

Tuckett, D., and Nikolic, M. (2017). The role of conviction and narrative in decision-making under radical uncertainty. Theory Psychol. 27, 501–523.

von Neumann, J., and Morgenstern, O. (1944). Theory of Games and Economic Behavior. Princeton University Press.

Wiessner, P. W. (2014). Embers of society: Firelight talk among the Ju/’hoansi Bushmen. Proc. Natl. Acad. Sci. U. S. A. 111, 14027–14035.

Willems, R. M., Nastase, S. A., and Milivojevic, B. (2020). Narratives for Neuroscience. Trends Neurosci. 43, 271–273.

Woolrich, M. W., Jbabdi, S., Patenaude, B., Chappell, M., Makni, S., Behrens, T., et al. (2009). Bayesian analysis of neuroimaging data in FSL. Neuroimage 45, S173–86.

Yarkoni, T., Poldrack, R. A., Nichols, T. E., Van Essen, D. C., and Wager, T. D. (2011). Large-scale automated synthesis of human functional neuroimaging data. Nat. Methods 8, 665–670.

Zhang, Z., Wang, S., Good, M., Hristova, S., Kayser, A. S., and Hsu, M. (2021). Retrieval-constrained valuation: Toward prediction of open-ended decisions. Proc. Natl. Acad. Sci. U. S. A. 118. doi: 10.1073/pnas.2022685118.

