## Supplementary material for "Neural support for contributions of utility and narrative processing of evidence in juror decision making": Figure 1-1

**Figure 1-1. Scenario crime descriptions and evidence options.**

| **Scenario** |  | | |
| --- | --- | --- | --- |
| **1** | **Crime Description** | Chelsea McNeil is accused of drug possession. McNeil allegedly sold two 80-microgram hits of LSD from an Altoids tin. | |
|  | **Criminal History** | **Related** | McNeil has a prior conviction for possession of methamphetamine. |
|  |  | **Unrelated** | McNeil has a prior conviction for reckless driving. |
|  |  | **No prior** | McNeil has no criminal record. |
|  | **Witness** | **Witness** | A witness saw McNeil accept cash in exchange for the tin's contents. |
|  |  | **No witness** | No witnesses saw McNeil with the LSD. |
|  | **Physical Evidence** | **DNA** | Hairs matching McNeil's DNA were found in a tin containing a small vial of LSD. |
|  |  | **Non DNA** | Fingerprints matching McNeil's were found on tin containing a small vial of LSD. |
|  |  | **None** | No physical evidence links McNeil to the drug-laden tin. |
| **2** | **Crime Description** | Jackie Archer stands accused of burglary. Archer allegedly entered a cafe after closing and took $230 from the cash register. | |
|  | **Criminal History** | **Related** | Archer was previously convicted of breaking and entering. |
|  |  | **Unrelated** | Archer was previously convicted of disturbing the peace. |
|  |  | **No prior** | Archer has no criminal record. |
|  | **Witness** | **Witness** | An eyewitness saw Archer enter the back door of the coffee shop after closing. |
|  |  | **No witness** | No witnesses saw Archer enter the coffee shop |
|  | **Physical Evidence** | **DNA** | Skin flakes matching Archer's DNA were found in the cash register. |
|  |  | **Non DNA** | A fingerprint matching Archer's was found on the cash register. |
|  |  | **None** | No physical evidence links Archer to the burglary. |
| **3** | **Crime Description** | Dan Montes is accused of theft. Montes allegedly took an unattended laptop from a library desk. | |
|  | **Criminal History** | **Related** | Montes was previously convicted of burglary. |
|  |  | **Unrelated** | Montes was previously convicted of driving while intoxicated. |
|  |  | **No prior** | Montes has no criminal record. |
|  | **Witness** | **Witness** | A witness saw Montes take the computer. |
|  |  | **No witness** | There were no witnesses to the laptop theft. |
|  | **Physical Evidence** | **DNA** | Police discovered hairs on the library desk matching Montes's DNA. |
|  |  | **Non DNA** | Cloth fibers matching a tear in Montes's jacket were found on the library desk. |
|  |  | **None** | No physical evidence links Dan to the theft. |
| **4** | **Crime Description** | Keith Galloway is on trial for bribing a public official. Galloway allegedly offered jewelry cash and other perks to a zoning official due to vote on a potential business development. | |
|  | **Criminal History** | **Related** | Galloway was previously convicted of making illegal campaign contributions. |
|  |  | **Unrelated** | Galloway was previously convicted of cocaine possession. |
|  |  | **No prior** | Galloway has no criminal record. |
|  | **Witness** | **Witness** | A golf caddy saw Galloway pass several pieces of jewelry to the official while playing on the course. |
|  |  | **No witness** | No one saw Galloway bribe the official. |
|  | **Physical Evidence** | **DNA** | Saliva matching Galloway's DNA was found on the seal of cash envelopes in the official's office. |
|  |  | **Non DNA** | Fingerprints matching Galloway's were found on cash envelopes at the official's office. |
|  |  | **None** | No physical evidence supports the bribery allegations. |
| **5** | **Crime Description** | Dennis Fray is accused of producing child pornography. Fray allegedly filmed 70 minutes of pornographic footage involving a child. | |
|  | **Criminal History** | **Related** | Fray was previously convicted of statutory rape. |
|  |  | **Unrelated** | Fray was previously convicted of tax fraud. |
|  |  | **No prior** | Fray has no criminal record. |
|  | **Witness** | **Witness** | An eyewitness testified that she saw Fray leading children into the hotel in which the videos were filmed. |
|  |  | **No witness** | There are no witnesses linking Fray to the pornographic videos. |
|  | **Physical Evidence** | **DNA** | Police discovered traces of semen matching Fray's DNA on memory sticks containing the footage. |
|  |  | **Non DNA** | Police discovered fingerprints matching Fray's on memory sticks containing the footage. |
|  |  | **None** | No physical evidence links Fray to the videos. |
| **6** | **Crime Description** | Quentin MacDonough is accused of shoplifting. MacDonough allegedly smuggled a guitar out of a music store in a case he carried in with him. | |
|  | **Criminal History** | **Related** | MacDonough was previously convicted of larceny. |
|  |  | **Unrelated** | MacDonough was previously convicted of reckless driving. |
|  |  | **No prior** | MacDonough has no prior convictions. |
|  | **Witness** | **Witness** | A witness testified that he saw MacDonough place the guitar into a case. |
|  |  | **No witness** | There were no witnesses to the theft. |
|  | **Physical Evidence** | **DNA** | Skin flakes matching MacDonough's DNA were found on the strings and bridge of the stolen guitar. |
|  |  | **Non DNA** | Fingerprints matching MacDonough's were found on the stolen guitar. |
|  |  | **None** | No physical evidence links MacDonough to the stolen guitar. |
| **7** | **Crime Description** | Dylan Turner stands accused of voluntary manslaughter. Turner allegedly bludgeoned his wife Marie and her lover with a table leg upon discovering them together in a motel killing them. | |
|  | **Criminal History** | **Related** | Turner was previously convicted of domestic assault. |
|  |  | **Unrelated** | Turner was previously convicted of income tax evasion. |
|  |  | **No prior** | Turner has no criminal history. |
|  | **Witness** | **Witness** | A witness saw Turner fleeing the hotel shortly after the killings. |
|  |  | **No witness** | There are no witnesses to link Turner to the killings. |
|  | **Physical Evidence** | **DNA** | Blood matching Dylan Turner's DNA was discovered in the hotel room. |
|  |  | **Non DNA** | Fingerprints matching Turner's were found on the table leg used to kill Marie. |
|  |  | **None** | No physical evidence links Turner to the killings. |
| **8** | **Crime Description** | | Teri Anderson is accused of telemarketing fraud. Anderson allegedly used a prepaid cell phone to make calls in which she impersonated an insurance agent to collect the personal information of two dozen victims. |
|  | **Criminal History** | **Related** | Anderson was previously convicted of wire fraud. |
|  |  | **Unrelated** | Anderson was previously convicted of driving while impaired. |
|  |  | **No prior** | Anderson has no record of prior offenses. |
|  | **Witness** | **Witness** | A store clerk testified that he sold Anderson the cell phone used to make the calls. |
|  |  | **No witness** | There are no witnesses linking Anderson to the calls. |
|  | **Physical Evidence** | **DNA** | Police found skin flakes matching Anderson's DNA in the casing of the phone used to place the calls. |
|  |  | **Non DNA** | Police found fingerprints matching Anderson's on the phone used to place the calls. |
|  |  | **None** | No physical evidence links Anderson to the phone used to place the calls. |
| **9** | **Crime Description** | Miranda Rasmussen is on trial for money laundering. Rasmussen an accountant allegedly helped a pair of convicted drug dealers disguise their profits by preparing financial documents on their behalf. | |
|  | **Criminal History** | **Related** | Rasmussen was previously convicted of extortion. |
|  |  | **Unrelated** | Rasmussen was previously convicted of providing alcohol to a minor. |
|  |  | **No prior** | Rasmussen has no record of any prior convictions. |
|  | **Witness** | **Witness** | A bystander saw the drug dealers entering and leaving Rasmussen's office. |
|  |  | **No witness** | No witnesses link Rasmussen to the drug dealers. |
|  | **Physical Evidence** | **DNA** | Hairs matching Rasmussen's DNA were found on financial documents seized from the drug dealers. |
|  |  | **Non DNA** | Fingerprints matching Rasmussen's were found on financial documents seized from the drug dealers. |
|  |  | **None** | No physical evidence links Rasmussen to the fraudulent transactions. |
| **10** | **Crime Description** | Maxine Franks is accused of first-degree murder. Franks allegedly waited in a parking deck for Gerald Dodd before fatally stabbing him in the back arm and throat with a six-inch knife. | |
|  | **Criminal History** | **Related** | Franks was previously convicted of assault. |
|  |  | **Unrelated** | Franks was previously convicted of writing bad checks. |
|  |  | **No prior** | Franks has no record of prior convictions. |
|  | **Witness** | **Witness** | A woman saw Franks enter the parking lot clutching a small bundle half an hour before Dodd's death. |
|  |  | **No witness** | No witnesses have linked Franks to Dodd's death. |
|  | **Physical Evidence** | **DNA** | Hairs matching Franks's DNA were found on and around Dodd's body. |
|  |  | **Non DNA** | Fingerprints matching Franks's were found on the handle of the knife used to kill Dodd. |
|  |  | **None** | No physical evidence links Franks to Dodd's death. |
| **11** | **Crime Description** | Luke Friedman is accused of second-degree murder. Friedman allegedly struck James Cole with a tire iron during a traffic altercation killing him. | |
|  | **Criminal History** | **Related** | Friedman was previously convicted of aggravated assault. |
|  |  | **Unrelated** | Friedman was previously convicted of possessing drug paraphernalia. |
|  |  | **No prior** | Friedman has no record of prior convictions. |
|  | **Witness** | **Witness** | A witness testified that she saw Friedman reaching into his trunk while arguing with Cole. |
|  |  | **No witness** | There were no witnesses to Cole's killing. |
|  | **Physical Evidence** | **DNA** | Blood matching Friedman's DNA was found on a sharp edge of the tire iron used to kill Cole. |
|  |  | **Non DNA** | Fingerprints matching Friedman's were found on the tire iron used to kill Cole. |
|  |  | **None** | No physical evidence links Friedman to Cole's death. |
| **12** | **Crime Description** | Steven Douglas stands accused of robbery. Douglas allegedly stole Sabine Hoffman's purse and $250 by approaching her from behind and claiming to have a gun. | |
|  | **Criminal History** | **Related** | Douglas was previously convicted of larceny. |
|  |  | **Unrelated** | Douglas was previously convicted of animal cruelty. |
|  |  | **No prior** | Douglas has no history of prior convictions. |
|  | **Witness** | **Witness** | A bystander identified Douglas as the only man hurrying from the scene. |
|  |  | **No witness** | There were no witnesses to the robbery. |
|  | **Physical Evidence** | **DNA** | Hoffman's empty purse was recovered nearby containing skin flakes and hair matching Douglas's DNA. |
|  |  | **Non DNA** | Hoffman's empty purse was recovered nearby bearing fingerprints matching Douglas's. |
|  |  | **None** | No physical evidence links Douglas to the robbery. |
| **13** | **Crime Description** | Harold Butler stands accused of sexual battery. Butler allegedly groped Carrie Bryant's thigh and buttocks while she was jogging. | |
|  | **Criminal History** | **Related** | Butler was previously convicted of rape. |
|  |  | **Unrelated** | Butler was previously convicted of reselling prescription drugs. |
|  |  | **No prior** | Butler has no criminal record. |
|  | **Witness** | **Witness** | A witness saw Butler grope Bryant. |
|  |  | **No witness** | There were no witnesses to the groping. |
|  | **Physical Evidence** | **DNA** | Skin flakes matching Butler's DNA were recovered from the fabric of Bryant's running shorts. |
|  |  | **Non DNA** | Police found a swatch of cloth matching a tear in Bryant's shorts in Butler's home. |
|  |  | **None** | No physical evidence supports the allegations against Butler. |
| **14** | **Crime Description** | | Miranda Hamlin stands accused of vehicular manslaughter. Hamlin allegedly crashed a stolen vehicle into Diana Masters's Prius killing her before fleeing the scene. |
|  | **Criminal History** | **Related** | Hamlin was previously convicted of reckless driving. |
|  |  | **Unrelated** | Hamlin was previously convicted of bribing a public official. |
|  |  | **No prior** | Hamlin has no criminal record. |
|  | **Witness** | **Witness** | A driver testified that Hamlin was the woman he saw running from the stolen vehicle. |
|  |  | **No witness** | There were no witnesses to the crash. |
|  | **Physical Evidence** | **DNA** | Police discovered blood matching Hamlin's on the steering wheel of the stolen vehicle. |
|  |  | **Non DNA** | Police discovered fingerprints matching Hamlin's on the steering wheel and gearshift of the stolen vehicle. |
|  |  | **None** | No physical evidence links Hamlin to the fatal crash. |
| **15** | **Crime Description** | | Virginia Davis is accused of stealing dogs from backyards and demanding a ransom from the owners for their safe return. |
|  | **Criminal History** | **Related** | Davis was previously convicted of animal cruelty. |
|  |  | **Unrelated** | Davis was previously convicted of running an unlicensed gambling operation. |
|  |  | **No prior** | Davis has no criminal history. |
|  | **Witness** | **Witness** | A postal carrier identified Davis as the woman he saw herding dogs into a van in the neighborhood. |
|  |  | **No witness** | There are no witnesses. |
|  | **Physical Evidence** | **DNA** | Police found skin flakes matching Davis's DNA on the post of an enclosure from which one of the dogs went missing. |
|  |  | **Non DNA** | Police have found Davis' fingerprints on the gates of several yards from which dogs went missing. |
|  |  | **None** | There is no physical evidence associating Davis with the thefts. |
| **16** | **Crime Description** | Atlee Newman is accused of shooting Melanie Farris while stealing her car. Newman allegedly dragged Farris out of her Acura before shooting her twice in the head and driving off. | |
|  | **Criminal History** | **Related** | Newman was previously convicted of mugging a couple at gunpoint. |
|  |  | **Unrelated** | Newman was previously convicted of poaching endangered bears. |
|  |  | **No prior** | Newman has no prior convictions. |
|  | **Witness** | **Witness** | A cyclist identified Newman as the driver of the Acura that nearly ran him off the road near the crime. |
|  |  | **No witness** | No witnesses observed Newman shoot Farris or steal the car. |
|  | **Physical Evidence** | **DNA** | Hairs matching Newman's DNA were found in the fabric of the Acura's driver's seat. |
|  |  | **Non DNA** | Newman's fingerprints were found on the Acura's steering wheel and dash. |
|  |  | **None** | No physical evidence links Newman to the carjacking. |
| **17** | **Crime Description** | Thomas Mitchell is accused of injuring two gang members in a drive-by shooting. | |
|  | **Criminal History** | **Related** | Mitchell previously served a sentence for assaulting a gang member with a knife. |
|  |  | **Unrelated** | Mitchell previously served a sentence for identity theft. |
|  |  | **No prior** | Mitchell has no prior convictions. |
|  | **Witness** | **Witness** | A witness saw Mitchell's red Suzuki and Mitchell's distinctive arm tattoos during the shooting. |
|  |  | **No witness** | There are no other witnesses. |
|  | **Physical Evidence** | **DNA** | Blood matching Mitchell's DNA was discovered on a handkerchief discarded near the scene. |
|  |  | **Non DNA** | Mitchell's fingerprints were discovered on the ejected bullet casings at the scene. |
|  |  | **None** | No physical evidence links Mitchell to the shooting. |
| **18** | **Crime Description** | Arthur Jefferson is accused of starting a four-alarm fire that burned his own warehouse to the ground in order to collect the insurance on it. | |
|  | **Criminal History** | **Related** | Jefferson was previously convicted of arson. |
|  |  | **Unrelated** | Jefferson was previously convicted of domestic assault. |
|  |  | **No prior** | Jefferson has no record of prior convictions. |
|  | **Witness** | **Witness** | An eyewitness saw Jefferson's red pickup pulling away just after the fire started. |
|  |  | **No witness** | There were no witnesses to who started the fire. |
|  | **Physical Evidence** | **DNA** | Traces of Jefferson's DNA were found on a lighter near the blaze. |
|  |  | **Non DNA** | Jefferson's fingerprints were found on a lighter near the blaze. |
|  |  | **None** | No physical evidence links Jefferson to the blaze. |
| **19** | **Crime Description** | Samuel Minor is accused of picking the lock on the front door of Thomas and Linda Harringtons' home as they slept and stealing a television a computer and some artwork. | |
|  | **Criminal History** | **Related** | Minor was previously convicted of breaking into a parked car to steal stereo equipment. |
|  |  | **Unrelated** | Minor was previously convicted of driving while impaired. |
|  |  | **No prior** | Minor has no record of previous criminal activity. |
|  | **Witness** | **Witness** | A neighbor testified that he saw Minor hunched over the doorknob of the front door for a long time before entering. |
|  |  | **No witness** | There were no witnesses to the break-in at the Harringtons' home. |
|  | **Physical Evidence** | **DNA** | Skin flakes containing Minor's DNA have been found near the broken lock of the Harringtons' front door. |
|  |  | **Non DNA** | Minor's fingerprints have been found along the doorjamb of the house. |
|  |  | **None** | No physical evidence links Minor to the scene. |
| **20** | **Crime Description** | | Mark Pearson is accused of locking 24-year-old Brianne Mills an African American student into the trunk of her car and pushing it into a river nearly drowning her. He also spray-painted the car with a large swastika and racial and sexual epithets including “slut,” “dyke,” and “nigger”. |
|  | **Criminal History** | **Related** | Pearson was previously convicted of battery alongside several known members of a local skinhead gang. |
|  |  | **Unrelated** | Pearson was previously convicted of using false identification to purchase alcohol. |
|  |  | **No prior** | Pearson has no record of prior convictions. |
|  | **Witness** | **Witness** | A fisherman walking by the river identified Pearson as the person who discarded a ski mask near the riverbank. |
|  |  | **No witness** | There were no witnesses to Mills's assault. |
|  | **Physical Evidence** | **DNA** | Police discovered blood containing Pearson's DNA under Mills's fingernails. |
|  |  | **Non DNA** | Pearson's fingerprints were found on a can of spray paint dropped by the sinking car. |
|  |  | **None** | There is no physical evidence linking Pearson to the crime. |
| **21** | **Crime Description** | Ronald Parker is accused of breaking through a townhouse back door entering the bedroom of Tammy King and raping her at knife-point. | |
|  | **Criminal History** | **Related** | Parker has a previous conviction for rape. |
|  |  | **Unrelated** | Parker has a previous conviction for fraud. |
|  |  | **No prior** | Parker has never been arrested for a felony. |
|  | **Witness** | **Witness** | The victim did not see her attacker but a neighbor identified Parker as the man who ran out of the house. |
|  |  | **No witness** | The victim did not see her attacker and there are no other witnesses. |
|  | **Physical Evidence** | **DNA** | Blood with Parker's DNA is found on the knife blade. |
|  |  | **Non DNA** | Fingerprints matching Parker's are found at the scene. |
|  |  | **None** | No physical evidence links Parker to the crime scene. |
| **22** | **Crime Description** | Robert Campbell is accused of mugging Jack and Rosa Smith in a grocery store parking lot. | |
|  | **Criminal History** | **Related** | Campbell has served prison time in the past for armed robbery. |
|  |  | **Unrelated** | Campbell has served prison time in the past for possession of heroin. |
|  |  | **No prior** | Campbell has no record of prior offenses. |
|  | **Witness** | **Witness** | A greeter at the store identified Campbell as the Smiths' assailant. |
|  |  | **No witness** | There were no other witnesses to the crime. |
|  | **Physical Evidence** | **DNA** | A pistol was found discarded nearby with Campbell's DNA on the grip. |
|  |  | **Non DNA** | Jack's empty wallet was found discarded nearby bearing Campbell's fingerprints. |
|  |  | **None** | No physical evidence links Campbell to the Smiths' assault. |
| **23** | **Crime Description** | | Jerry Lee a known gang member is accused of the murder of a grocery store owner and his wife after they refused to pay him 'protection' money. |
|  | **Criminal History** | **Related** | Lee has a prior conviction for extortion. |
|  |  | **Unrelated** | Lee has a prior conviction for possession of crack cocaine. |
|  |  | **No prior** | Lee has no criminal record. |
|  | **Witness** | **Witness** | A neighboring shopkeeper identified Lee as the man fleeing after the shooting. |
|  |  | **No witness** | No witnesses saw Lee at the scene. |
|  | **Physical Evidence** | **DNA** | Blood matching Lee's DNA was found on a broken bottle near the grocer's body. |
|  |  | **Non DNA** | Lee's fingerprints were found on the pistol used in the shooting. |
|  |  | **None** | No physical evidence links Lee to the murders. |
| **24** | **Crime Description** | Christopher Pepper and Alva Bennett are accused of manufacturing counterfeit cashier's cheques using a printing press located in a condemned office building. | |
|  | **Criminal History** | **Related** | Pepper has a past conviction for fraud and Bennett for selling knockoff handbags on eBay. |
|  |  | **Unrelated** | Pepper has a past conviction for battery and Bennett for driving with a suspended license. |
|  |  | **No prior** | Neither Pepper nor Bennett has any prior record of convictions. |
|  | **Witness** | **Witness** | A barista at a nearby cafe observed Pepper and Bennett going in and out of the condemned building. |
|  |  | **No witness** | No witnesses observed Peppers or Bennett entering the condemned building during the past few months. |
|  | **Physical Evidence** | **DNA** | Hair and skin flakes with Pepper's and Bennett's DNA were found on and around the counterfeit press. |
|  |  | **Non DNA** | Pepper's and Bennett's fingerprints were found on and around the counterfeit press. |
|  |  | **None** | No physical evidence links Pepper or Bennett to the counterfeiting. |
| **25** | **Crime Description** | Janet Norment is accused of smuggling a suitcase of marijuana into the United States. | |
|  | **Criminal History** | **Related** | Norment was recently released from prison following a sentence for manufacturing methamphetamine. |
|  |  | **Unrelated** | Norment was recently released from prison following an armed robbery sentence. |
|  |  | **No prior** | Norment has no prior convictions. |
|  | **Witness** | **Witness** | A check-in attendant at the airport identified Norment as the woman who checked the suitcase in. |
|  |  | **No witness** | No witnesses saw Norment check the bags in. |
|  | **Physical Evidence** | **DNA** | Eyelashes with Norment's DNA were discovered in the suitcase. |
|  |  | **Non DNA** | Cloth fibers matching several articles of clothing in Norment's home were discovered in the suitcase. |
|  |  | **None** | No physical evidence was discovered linking Norment to the luggage. |
| **26** | **Crime Description** | Trevor Allen and Kenneth Green are accused of leaving 13 undocumented immigrants to die in a truck. Allen and Green were allegedly smuggling the immigrants across the U.S. border but after the truck stopped working they allegedly fled while leaving the immigrants locked in the back. By the time the immigrants were found they had all died of exposure. | |
|  | **Criminal History** | **Related** | Green and Allen both have prior convictions for illegally transporting noncitizens into the United States. |
|  |  | **Unrelated** | Green has a prior conviction for marijuana possession and Allen for domestic violence. |
|  |  | **No prior** | Neither Green nor Allen has a record of prior convictions. |
|  | **Witness** | **Witness** | A gas station attendant identified Green and Allen as the men who borrowed his phone to call for a ride near where the truck was discovered. |
|  |  | **No witness** | No witnesses saw either of the men load or depart from the vehicle. |
|  | **Physical Evidence** | **DNA** | Green's and Allen's DNA was found on the lips of several empty soda bottles in the truck. |
|  |  | **Non DNA** | Green and Allen's fingerprints were found on the truck's front seat. |
|  |  | **None** | No physical evidence links Allen or Green to the truck. |
| **27** | **Crime Description** | Clarence Warren is accused of operating an unlicensed distillery. Police discovered a makeshift pot still several hundred pounds of white sugar some coiled copper piping and forty gallons of moonshine on an abandoned farm. | |
|  | **Criminal History** | **Related** | Warren has a previous conviction for manufacturing methamphetamine. |
|  |  | **Unrelated** | Warren has a previous conviction for armed robbery. |
|  |  | **No prior** | Warren has no record of prior convictions. |
|  | **Witness** | **Witness** | An electrical maintenance worker testified that he saw Warren around the still. |
|  |  | **No witness** | No witnesses saw Warren in or around the farmhouse. |
|  | **Physical Evidence** | **DNA** | Warren's DNA was found on the lips of mason jars at the scene. |
|  |  | **Non DNA** | A strip of fabric matching a tear in Warren's jacket was found on a nail protruding from the farmhouse. |
|  |  | **None** | No physical evidence links Warren to the distillery. |
| **28** | **Crime Description** | Jeff Martinez is accused of bombing a shopping center. An unregistered black pickup truck was loaded with explosives driven through the main entrance and detonated after the driver fled. | |
|  | **Criminal History** | **Related** | Martinez was previously convicted of aggravated arson. |
|  |  | **Unrelated** | Martinez was previously convicted of indecent exposure. |
|  |  | **No prior** | Martinez has no criminal history. |
|  | **Witness** | **Witness** | A bystander identified Martinez as the truck's driver. |
|  |  | **No witness** | No witnesses were able to identify the truck's driver. |
|  | **Physical Evidence** | **DNA** | Blood matching Martinez's DNA was found on a piece of shrapnel near the mall entrance. |
|  |  | **Non DNA** | Martinez's fingerprints were found on an undamaged section of the truck. |
|  |  | **None** | No physical evidence links Martinez to the bombing. |
| **29** | **Crime Description** | Bret Botham is accused of critically injuring two people while illegally racing. The car collided with a stopped van after he skidded out of control on a patch of gravel. | |
|  | **Criminal History** | **Related** | Botham had his license suspended last year for driving recklessly. |
|  |  | **Unrelated** | Botham had been convicted of shoplifting last year. |
|  |  | **No prior** | Botham has no prior convictions. |
|  | **Witness** | **Witness** | The driver of the van testified that Botham was the driver. |
|  |  | **No witness** | The driver of the van dazed by the impact was unable to identify the fleeing motorist. |
|  | **Physical Evidence** | **DNA** | The car's driver fled into the woods leaving blood with Botham's DNA on the car's airbag and dash. |
|  |  | **Non DNA** | The car's driver fled into the woods but Botham's fingerprints were found on the car's dash console and gearshift. |
|  |  | **None** | The car's driver fled into the woods leaving no recoverable physical evidence. |
| **30** | **Crime Description** | Virginia MacPherson is accused of sending anthrax-laced envelopes to a United States senator and the father of her boyfriend. After intercepting the envelopes the FBI arrested MacPherson. | |
|  | **Criminal History** | **Related** | As a high school student MacPherson sent threatening letters to the parents of a previous boyfriend. |
|  |  | **Unrelated** | MacPherson had previously been convicted for shoplifting. |
|  |  | **No prior** | MacPherson has no criminal record. |
|  | **Witness** | **Witness** | A researcher testified that he saw MacPherson walking around a lab from which anthrax samples went missing. |
|  |  | **No witness** | Investigators were unable to find anyone who could link MacPherson to the anthrax. |
|  | **Physical Evidence** | **DNA** | FBI agents discovered saliva matching MacPherson's DNA on the stamps of several envelopes. |
|  |  | **Non DNA** | FBI agents discovered MacPherson's fingerprints on several of the tainted envelopes. |
|  |  | **None** | No physical evidence was found to link MacPherson to the envelopes. |
| **31** | **Crime Description** | Nathan Lewis is accused of sexually abusing a fourteen-year-old boy after police find a camera with pornographic images of the boy in the junior high locker room. | |
|  | **Criminal History** | **Related** | Lewis was previously convicted of sexually harassing an intern. |
|  |  | **Unrelated** | Lewis was previously convicted of fraud. |
|  |  | **No prior** | Lewis has no prior convictions. |
|  | **Witness** | **Witness** | The boy is too traumatized to testify but a school custodian saw Saunders carrying photo equipment into the locker room. |
|  |  | **No witness** | The boy is too traumatized to testify and there are no witnesses. |
|  | **Physical Evidence** | **DNA** | Semen with Lewis's DNA was found on the camera. |
|  |  | **Non DNA** | Lewis's fingerprints were found on the camera. |
|  |  | **None** | No physical evidence links Lewis to the photos. |
| **32** | **Crime Description** | Brian Jones is accused of shooting Judge Timothy Baker in the back outside a courthouse. As Jones fled he banged his hand into a door jamb and dropped his gun. | |
|  | **Criminal History** | **Related** | Jones was previously convicted of battery. |
|  |  | **Unrelated** | Jones was previously convicted of fraud. |
|  |  | **No prior** | Jones has no criminal record. |
|  | **Witness** | **Witness** | A bystander saw Jones running away after she heard a gunshot. |
|  |  | **No witness** | No witnesses were present in the courthouse alley to identify Baker's attacker. |
|  | **Physical Evidence** | **DNA** | Blood with Jones's DNA is found on an ejected bullet casing. |
|  |  | **Non DNA** | Jones's fingerprints were found on the pistol's grip and trigger. |
|  |  | **None** | No usable physical evidence identifying Jones could be found. |
| **33** | **Crime Description** | Samantha Harris is accused of vehicular manslaughter during a hit-and-run. She had been texting while driving and collided with Cameron Bailey who later died. | |
|  | **Criminal History** | **Related** | Harris had previously been convicted of leaving a child she was babysitting locked in a hot car. |
|  |  | **Unrelated** | Harris had previously been convicted of shoplifting. |
|  |  | **No prior** | Harris has no criminal record. |
|  | **Witness** | **Witness** | A witness saw Harris texting at a red light two blocks from where the accident happened. |
|  |  | **No witness** | No witness saw Harris either texting or driving. |
|  | **Physical Evidence** | **DNA** | A paper soda cup with Harris's saliva was found near Bailey's body. |
|  |  | **Non DNA** | Paint matching the color of Harris's car was found scraped onto Bailey's belt buckle. |
|  |  | **None** | There was no physical evidence found linking Harris to Bailey. |
