## Supplementary material for "Neural support for contributions of utility and narrative processing of evidence in juror decision making": Figure 4-1

**Extended Data**

**Figure 4-1. Neurosynth topic maps included in cognitive models and submodels.**

| **Model/Submodel** | | | | **Topic** | **Top Terms** |
| --- | --- | --- | --- | --- | --- |
| Social-Affective | Affect | | | [126](https://neurosynth.org/analyses/topics/v5-topics-200/126) | anxiety, aggression, anxious, trait, amygdala, cr, disorder, gad, anticipation, emotional, anticipatory, high, aggressive, ha, social, worry, fear, temperament, individuals, panic, scores, anger, threat, symptoms, phobia, reactivity, dental, relationship, limbic, iu, provocation, reactive, generalized, characterized, subclinical, bi, regulation, insula, somatic, paradigm |
|  |  |  |  | [139](https://neurosynth.org/analyses/topics/v5-topics-200/139) | faces, emotional, facial, expressions, emotion, amygdala, neutral, fearful, happy, face, sad, emotions, angry, expression, social, fear, response, perception, affective, recognition, anger, affect, disgust, viewing, happiness, gender, sadness, magnetic, fusiform, presented, resonance, interaction, emotionally, salient, expressing, masked, threat, limbic, viewed, sensitivity |
|  |  |  |  | [180](https://neurosynth.org/analyses/topics/v5-topics-200/180) | amygdala, threat, fear, anxiety, response, avoidance, reactivity, threatening, habituation, responding, shock, aversive, heightened, emotional, cues, approach, affective, amygdalar, threats, bnst, research, bla, individuals, emotion, basolateral, hippocampus, danger, periaqueductal, limbic, nucleus, amygdalae, implicated, defensive, stress, insula, trait, exaggerated, safe, anxious, startle |
|  | Moral Judgment | Social | | [145](https://neurosynth.org/analyses/topics/v5-topics-200/145) | mind, mental, social, tom, states, theory, mentalizing, belief, state, attribution, person, reasoning, empathy, people, cognitive, beliefs, affective, junction, medial, prefrontal, precuneus, perspective, inference, intentional, inferences, intentions, wandering, lsf, cognition, inferring, agency, temporoparietal, stories, mpfc, infer, physical, recruited, hsf, temporo, emotions |
|  |  |  |  | [154](https://neurosynth.org/analyses/topics/v5-topics-200/154) | social, interactions, interaction, cognition, partner, human, interpersonal, exclusion, person, oxytocin, people, game, socially, cooperation, partners, individuals, ot, rejection, trust, participant, insula, played, mentalizing, context, relationships, nonsocial, animacy, cues, cooperative, interactive, friends, paradigm, predicted, friend, prosocial, correlates, political, animate, personal, ball |
|  |  | Moral | | [112](https://neurosynth.org/analyses/topics/v5-topics-200/112) | context, empathy, contextual, contexts, ambiguity, ambiguous, empathic, dominant, prosocial, concern, modulated, resolution, situations, automatic, unambiguous, presence, subordinate, meanings, contextually, altruistic, frequent, social, representations, interpretation, flexible, embedded, situational, affects, empathize, scanned, donation, architecture, disambiguating, embedding, settings, occurs, dominance, oriented, occurred, involve |
|  |  |  |  | [135](https://neurosynth.org/analyses/topics/v5-topics-200/135) | basal, ganglia, moral, guilt, harm, judgments, judgment, thalamus, forebrain, care, transgressions, scenarios, violations, gratitude, actions, lying, harmful, morally, justice, personal, wrong, dilemmas, acts, judged, cost, sentiments, cholinergic, benefit, bad, morality, norms, issues, conventional, good, utilitarian, anti, accidental, unified, legal, shame |
| Evidentiary | Utility | Probability | | [002](https://neurosynth.org/analyses/topics/v5-topics-200/002) | problem, problems, arithmetic, solving, calculation, mental, mathematical, math, ag, addition, angular, multiplication, parietal, subtraction, operations, retrieval, operation, number, solution, gyrus, ef, solved, sulcus, numerical, insight, solutions, intraparietal, competence, solve, single, strategies, adults, cognitive, cognition, mathematics, arithmetical, small, simple, digit, size |
|  |  |  |  | [086](https://neurosynth.org/analyses/topics/v5-topics-200/086) | ips, number, numerical, intraparietal, numbers, magnitude, sulcus, symbolic, parietal, distance, dd, representation, numerosity, digit, small, digits, size, large, arabic, estimation, quantifiers, nonsymbolic, magnitudes, quantity, counting, larger, arrays, quantities, abstract, symbols, line, letters, dyscalculia, congruity, sulci, dots, notation, fractions, numerals, adults |
|  |  |  |  | [128](https://neurosynth.org/analyses/topics/v5-topics-200/128) | prediction, error, outcome, errors, predictive, probability, expectation, predictions, outcomes, events, expectations, uncertainty, predictability, unexpected, predicted, predict, model, environment, unpredictable, models, predictable, expectancy, entropy, reflected, future, cancer, chemotherapy, predicting, likelihood, adjustment, probabilities, pe, updating, prior, coding, adaptation, upcoming, omission, theories, environmental |
|  |  | Value | | [010](https://neurosynth.org/analyses/topics/v5-topics-200/010) | ofc, orbitofrontal, cortex, lateral, medial, amygdala, mofc, frontal, evaluation, frontopolar, pole, prefrontal, limbic, systems, orbital, regional, lofc, paralimbic, reward, negative, specifically, orbito, serving, associations, plays, values, functioning, immediately, unfavorable, sectors, social, successful, olfactocentric, vigilant, favorable, sections, declared, reconfiguration, temporopolar, bmrmi |
|  |  |  |  | [186](https://neurosynth.org/analyses/topics/v5-topics-200/186) | decision, making, choice, decisions, choices, uncertainty, outcomes, discounting, outcome, striatum, rewards, options, valuation, preferences, make, values, preference, probability, reward, subjective, individual, gains, economic, losses, loss, option, process, choose, aversion, gambling, insula, reputation, risky, selection, intertemporal, accumulation, people, prior, igt, choosing |
|  |  |  |  | [197](https://neurosynth.org/analyses/topics/v5-topics-200/197) | reward, striatum, anticipation, ventral, monetary, rewards, motivation, loss, response, incentive, striatal, motivational, sensitivity, punishment, gain, outcomes, outcome, rewarding, money, delay, magnitude, mesolimbic, losses, gains, anticipatory, midbrain, salience, receipt, aversive, gambling, motivated, appetitive, rewarded, cues, feedback, incentives, dopaminergic, win, anticipated, avoidance |
|  | Narrative | Experience, Culture, & Recall | Culture & Ideation Bias | [030](https://neurosynth.org/analyses/topics/v5-topics-200/030) | events, future, personal, past, cultural, thinking, construction, episodic, experiences, imagining, culture, imagined, counterfactual, autobiographical, engaged, simulation, american, experience, imagination, hippocampus, simulations, construal, world, real, remembering, engage, contextual, east, thought, elaboration, cultures, projection, imagine, interdependent, statements, thoughts, core, ep, western, episodes |
|  |  |  |  | [100](https://neurosynth.org/analyses/topics/v5-topics-200/100) | gestures, abstract, race, concrete, intention, gesture, communicative, prospective, intentions, speech, racial, communication, context, actor, iconic, stereotypes, mp, indirect, pointing, american, black, action, videos, stereotype, unrelated, white, aa, intended, ig, performing, verbal, meaningless, isolated, prejudice, gr, situation, biases, chinese, ongoing, img |
|  |  |  | Recall | [028](https://neurosynth.org/analyses/topics/v5-topics-200/028) | memory, retrieval, episodic, recollection, memories, autobiographical, recall, hippocampus, events, recognition, familiarity, semantic, hippocampal, context, retrieved, lateral, reactivation, recency, suggest, temporal, medial, source, remember, parietal, remembering, remote, past, item, retrieve, engaged, successful, supporting, parahippocampal, correlates, encoding, posterior, monitoring, ams, precuneus, reinstatement |
|  |  |  |  | [111](https://neurosynth.org/analyses/topics/v5-topics-200/111) | memory, encoding, hippocampal, hippocampus, retrieval, subsequent, successful, episodic, recognition, formation, encoded, recall, remembered, associative, items, associations, performance, binding, verbal, memories, success, learning, subsequently, suggest, predicted, correlates, phase, parahippocampal, prefrontal, anterior, forgotten, tested, test, relational, versus, paired, recognized, declarative, scanning, remember |
|  |  | Reading | | [009](https://neurosynth.org/analyses/topics/v5-topics-200/009) | semantic, word, knowledge, words, temporal, meaning, retrieval, concepts, atl, lexical, semantically, anterior, inferior, unrelated, pairs, lifg, association, language, demands, abstract, phonological, decision, relatedness, concept, correlates, semantics, imageability, controlled, picture, names, memory, associations, judgments, lexico, concrete, meanings, required, nouns, selection, level |
|  |  |  |  | [093](https://neurosynth.org/analyses/topics/v5-topics-200/093) | language, sentences, comprehension, sentence, syntactic, linguistic, pars, semantic, broca, temporal, meaning, literal, narrative, posterior, opercularis, inferior, discourse, metaphors, story, triangularis, spoken, syntax, production, read, reading, speech, listening, metaphor, text, lifg, word, hemisphere, stories, presented, pragmatic, context, german, structure, listened, irony |
|  |  |  |  | [095](https://neurosynth.org/analyses/topics/v5-topics-200/095) | verbs, verb, nouns, noun, grammatical, language, structure, generation, production, subject, lexical, morphological, phrases, word, regular, argument, words, syntactic, object, inferior, linguistic, number, initial, frontal, ft, sentence, np, irregular, violations, forms, thematic, lifg, tense, morphology, english, inflected, lmtg, past, types, sentences |
|  |  |  |  | [162](https://neurosynth.org/analyses/topics/v5-topics-200/162) | words, word, lexical, frequency, recognition, visual, semantic, presented, form, vwfa, reading, pseudowords, decision, posterior, meaning, high, written, phonological, pseudo, spoken, linguistic, read, real, lists, nonwords, involvement, sublexical, visually, representations, number, competition, length, engaged, material, forms, neighborhood, silent, highly, syllable, strings |
|  |  | Working Memory | | [020](https://neurosynth.org/analyses/topics/v5-topics-200/020) | wm, load, memory, working, task, maintenance, performance, ltm, high, capacity, prefrontal, distraction, increasing, verbal, loads, probe, parietal, demands, maintained, delay, demand, manipulation, accuracy, levels, resources, maintain, held, parametric, paradigm, eos, recruitment, susceptibility, encoding, sternberg, sample, distracters, probes, letter, secondary, memoranda |
|  |  |  |  | [179](https://neurosynth.org/analyses/topics/v5-topics-200/179) | memory, working, task, verbal, maintenance, performance, load, cognitive, storage, updating, capacity, rehearsal, spatial, network, function, executive, functions, manipulation, probe, phase, visuospatial, frontoparietal, swm, phonological, high, sternberg, retention, mnemonic, maintain, articulatory, delayed, items, held, wmc, buffer, maintaining, store, remember, regional, operations |
