## Supplementary material for "Neural support for contributions of utility and narrative processing of evidence in juror decision making": Figure 4-2

**Extended Data**

**Figure 4-2. Spatial similarity (linear regression fit) for juror decision-making models and thresholded brain activation patterns associated with evidence accumulation and crime-type bias.** For evidence accumulation and crime-type bias fMRI maps, the table includes overall model fit (BIC score), p-value, and topic map standardized coefficient measures (beta and standard error (se)). P-values reflect the number of BIC values from the null distribution (n=5,000 permutations) that are greater than the model's BIC.

| **Neurosynth Decision Making Model** | **Model Fit (BIC)** | **Topic #** | **Coefficients** | |
| --- | --- | --- | --- | --- |
|  | **Evidence Accumulation** |  | **Evidence**  **Accumulation** | |
|  |  |  | **beta** | **se** |
| Social-Affective | BIC = 41947, p = 0.720 | intercept | 2.958 | 0.003 |
|  |  | 126 | -0.032 | 0.004 |
|  |  | 139 | -0.072 | 0.004 |
|  |  | 180 | 0.011 | 0.005 |
|  |  | 112 | 0.04 | 0.003 |
|  |  | 135 | -0.029 | 0.003 |
|  |  | 145 | -0.009 | 0.003 |
|  |  | 154 | -0.003 | 0.004 |
| Evidentiary | BIC = 39548, p = 0.002 | intercept | 2.958 | 0.003 |
|  |  | 2 | 0.071 | 0.004 |
|  |  | 10 | -0.012 | 0.003 |
|  |  | 86 | 0.012 | 0.004 |
|  |  | 128 | 0.005 | 0.004 |
|  |  | 186 | 0.04 | 0.004 |
|  |  | 197 | 0.027 | 0.004 |
|  |  | 28 | 0.047 | 0.004 |
|  |  | 30 | -0.014 | 0.003 |
|  |  | 100 | -0.022 | 0.003 |
|  |  | 111 | 0.026 | 0.004 |
|  |  | 9 | 0.108 | 0.005 |
|  |  | 93 | -0.02 | 0.005 |
|  |  | 95 | 0.026 | 0.005 |
|  |  | 162 | 0.004 | 0.004 |
|  |  | 20 | -0.019 | 0.004 |
|  |  | 179 | 0.013 | 0.004 |
| Narrative | BIC = 40285, p = 0.014 | intercept | 2.958 | 0.003 |
|  |  | 28 | 0.068 | 0.004 |
|  |  | 30 | -0.024 | 0.003 |
|  |  | 100 | -0.033 | 0.003 |
|  |  | 111 | 0.014 | 0.004 |
|  |  | 9 | 0.115 | 0.005 |
|  |  | 93 | -0.037 | 0.005 |
|  |  | 95 | 0.022 | 0.005 |
|  |  | 162 | 0.007 | 0.004 |
|  |  | 20 | -0.002 | 0.004 |
|  |  | 179 | 0.032 | 0.004 |
| Utility | BIC = 41550, p = 0.241 | intercept | 2.958 | 0.003 |
|  |  | 2 | 0.104 | 0.004 |
|  |  | 10 | -0.019 | 0.003 |
|  |  | 86 | -0.015 | 0.004 |
|  |  | 128 | -0.021 | 0.004 |
|  |  | 186 | 0.05 | 0.004 |
|  |  | 197 | 0.005 | 0.004 |
| Affect | BIC = 42145, p = 0.370 | intercept | 2.958 | 0.003 |
|  |  | 126 | -0.039 | 0.004 |
|  |  | 139 | -0.063 | 0.004 |
|  |  | 180 | 0.007 | 0.004 |
| Moral Judgment | BIC = 42524, p = 0.893 | intercept | 2.958 | 0.003 |
|  |  | 112 | 0.035 | 0.003 |
|  |  | 135 | -0.025 | 0.003 |
|  |  | 145 | -0.013 | 0.003 |
|  |  | 154 | -0.032 | 0.003 |
| Experience, Culture, & Recall Narrative | BIC = 41436, p = 0.065 | intercept | 2.958 | 0.003 |
|  |  | 28 | 0.094 | 0.004 |
|  |  | 30 | -0.045 | 0.003 |
|  |  | 100 | -0.015 | 0.003 |
|  |  | 111 | 0.05 | 0.003 |
| Reading Narrative | BIC = 40975, p = 0.020 | intercept | 2.958 | 0.003 |
|  |  | 9 | 0.147 | 0.005 |
|  |  | 93 | -0.048 | 0.005 |
|  |  | 95 | 0.025 | 0.005 |
|  |  | 162 | 0.001 | 0.004 |
| Working Memory Narrative | BIC = 42522, p = 0.578 | intercept | 2.958 | 0.003 |
|  |  | 20 | -0.01 | 0.004 |
|  |  | 179 | 0.06 | 0.004 |
| Utility (Probability) | BIC = 41756, p = 0.095 | intercept | 2.958 | 0.003 |
|  |  | 2 | 0.109 | 0.004 |
|  |  | 86 | -0.012 | 0.004 |
|  |  | 128 | -0.001 | 0.003 |
| Utility (Value) | BIC = 42372, p = 0.613 | intercept | 2.958 | 0.003 |
|  |  | 10 | -0.04 | 0.003 |
|  |  | 186 | 0.067 | 0.004 |
|  |  | 197 | -0.031 | 0.004 |
| Moral Judgment (Social) | BIC = 42684, p = 0.822 | intercept | 2.958 | 0.003 |
|  |  | 145 | -0.008 | 0.003 |
|  |  | 154 | -0.033 | 0.003 |
| Moral Judgment (Moral Judgment) | BIC = 42637, p = 0.739 | intercept | 2.958 | 0.003 |
|  |  | 112 | 0.026 | 0.003 |
|  |  | 135 | -0.034 | 0.003 |
| Experience, Culture, & Recall (Culture & Ideation Bias) | BIC = 42800, p=0.962 | intercept | 2.958 | 0.003 |
|  |  | 30 | -0.005 | 0.003 |
|  |  | 100 | -0.012 | 0.003 |
| Experience, Culture, & Recall (Recall) | BIC = 41636, p = 0.019 | intercept | 2.958 | 0.003 |
|  |  | 28 | 0.077 | 0.003 |
|  |  | 111 | 0.051 | 0.003 |
