## Supplementary material for "Neural support for contributions of utility and narrative processing of evidence in juror decision making": Figure 4-3

**Extended Data**

**Figure 4-3. Null model spatial similarity threshold for juror decision-making models and thresholded brain activation patterns associated with evidence accumulation.** For evidence accumulation and crime-type bias fMRI maps, the table includes overall model fit (BIC score) threshold dependent on the number of topics included in a model. The threshold BIC is the 95th percentile of the null distribution of 5,000 random models.

| **Model** | **Number of Topics** | **Similarity Threshold (BIC)** |
| --- | --- | --- |
|  |  | Evidence Accumulation |
| Moral Judgment (Social) | 2 | 41828 |
| Moral Judgment (Moral) |  |  |
| Experience, Culture, & Recall (Culture & Ideation Bias) |  |  |
| Experience, Culture, & Recall (Recall) |  |  |
| Working Memory |  |  |
| Affect | 3 | 41595 |
| Utility (Value) |  |  |
| Utility (Probability) |  |  |
| Moral Judgment | 4 | 41333 |
| Experience, Culture, & Recall |  |  |
| Reading |  |  |
| Utility | 6 | 41000 |
| Social-Affective | 7 | 40885 |
| Narrative | 10 | 40544 |
| Evidentiary | 16 | 40059 |
